# Nr4a1 regulates inhibitory circuit structure and function in the mouse brain

**DOI:** 10.1101/2022.06.14.496205

**Authors:** Min Huang, Simon Pieraut, Jasmine Cao, Filip de Souza Polli, Vincenzo Roncace, Gloria Shen, Anton Maximov

## Abstract

Central neurons express unique repertoires of secreted and transmembrane proteins that define their synaptic connectivity. However, how these molecular programs are regulated remains poorly understood. Our study shows that, in inhibitory GABAergic interneurons in the mouse forebrain, transcription of synaptic organizer molecules is controlled by Nr4a1, a nuclear receptor whose expression is transiently induced by sensory experience and required for normal learning. Nr4a1 exserts opposite effects on local axonal wiring of Parvalbumin- and Somatostatin-positive interneurons that innervate different subcellular domains of their postsynaptic partners. Loss of Nr4a1 activity in these interneurons leads to cell-type-specific transcriptional switches in multiple gene families, including those involved in surface adhesion and repulsion. Our findings reveal a mechanism by which inducible transcription factors dynamically alter the combinatorial synaptic organizing codes for structural plasticity.

---

Assembly of neural circuits in the developing brain is orchestrated by molecular programs that define the cellular and subcellular patterns of synaptic connectivity through combinatorial expression of multiple gene families (*1-3*). Over the course of postnatal lifespan, central neurons repetitively change their gene expression profiles in response to sensory experience. These events begin with transient induction of transcription factors (TFs) encoded by early response genes (ERGs, also referred to as immediate early genes) whose mRNAs and proteins are rapidly synthesized in cellular ensembles activated by excitatory synaptic input and calcium influx (*4*). Experimental evidence suggests that activity-regulated transcription is necessary for dynamic rewiring of the brain and memory storage (*4, 5*). However, the natures of ERG effectors in diverse neuron classes are poorly understood. In this work, we studied how ERG signaling impacts transcriptional programs underlying the architecture and function of inhibitory circuits.

In the mammalian cerebral cortex and hippocampus, synaptic inhibition is mediated by ∼20 subtypes of GABAergic interneurons (INs) with distinct morphologies, postsynaptic target specificities, and physiological properties (*6*). Among these INs, the two most abundant populations express Parvalbumin (PV) and Somatostatin (Sst). PV^+^ fast spiking basket cells innervate the somas of glutamatergic projection neurons (PNs), whereas Sst^+^ Martinotti cells provide GABAergic inputs onto distal parts of PN dendrites (*6*). Considering that PV^+^ and Sst^+^ cells are involved in memory coding (*7-9*), we searched for TFs which could be recruited in these INs during associative learning. We accomplished this task using RiboTag, a technique for immuno-isolation of translating mRNAs in complexes with ribosomes containing a genetically targeted HA epitope-tagged protein, Rpl22 (*10*). We introduced RiboTag into each IN population by crossing the Cre recombinase-dependent *Rpl22-HA* mouse allele with *Pvalb*^*Cre*^ and *Sst*^*Cre*^ drivers (*11, 12*), extracted cell-specific pools of mRNAs from the cortices and hippocampi of young adults at postnatal day (p) 60, and analyzed these mRNAs by deep sequencing (RNA-seq) (Fig. 1A). As the first step, we performed quality controls to ensure appropriate enrichment of markers of IN lineages and negligible contamination with markers of other cell classes in mRNA libraries from *Pvalb*^*Cre*^/*RiboTag* and *Sst*^*Cre*^/*RiboTag* mice (fig. S1). We then surveyed acute changes in IN transcriptomes that were elicited in whole animals by contextual fear conditioning (CFC), a neurobehavioral paradigm for acquisition of stable associative memory (*13*) (Fig. 1, B and C). In parallel experiments, *Pvalb*^*Cre*^/*RiboTag* and *Sst*^*Cre*^/*RiboTag* mice received single doses of the GABA receptor antagonist, pentylenetetrazol (PTZ) (*14*), to identify genes whose mRNA levels were altered by artificial widespread network excitation as a frame of reference (Fig. 1, D and E). We selected all known and putative TFs from sets of significantly up- and down-regulated genes with the false discovery rate (FDR) of <0.05, ranked them by fold-change (FC) of mRNA levels, and compared expression across datasets. These analyses prompted us to investigate Nr4a1, an orphan nuclear hormone receptor encoded by a stress-sensitive ERG (*15-17*) whose role in GABAergic INs remained unknown. Nr4a1 was strongly induced in PV^+^ (FC=7.3; FDR=2.2^-11^) and Sst^+^ (FC=5.4; FDR=4.2^-26^) cells of fear-conditioned mice (30 minutes post-CFC), and was the only TF activated in both groups of INs under two experimental settings. By contrast, CFC failed to trigger a similar increase in mRNAs of several other classical ERGs, including Npas4 whose transcription is regulated in cortical INs by visual experience (*18, 19*) (Fig. 1, F and G and fig. S2).

**Fig. 1.**
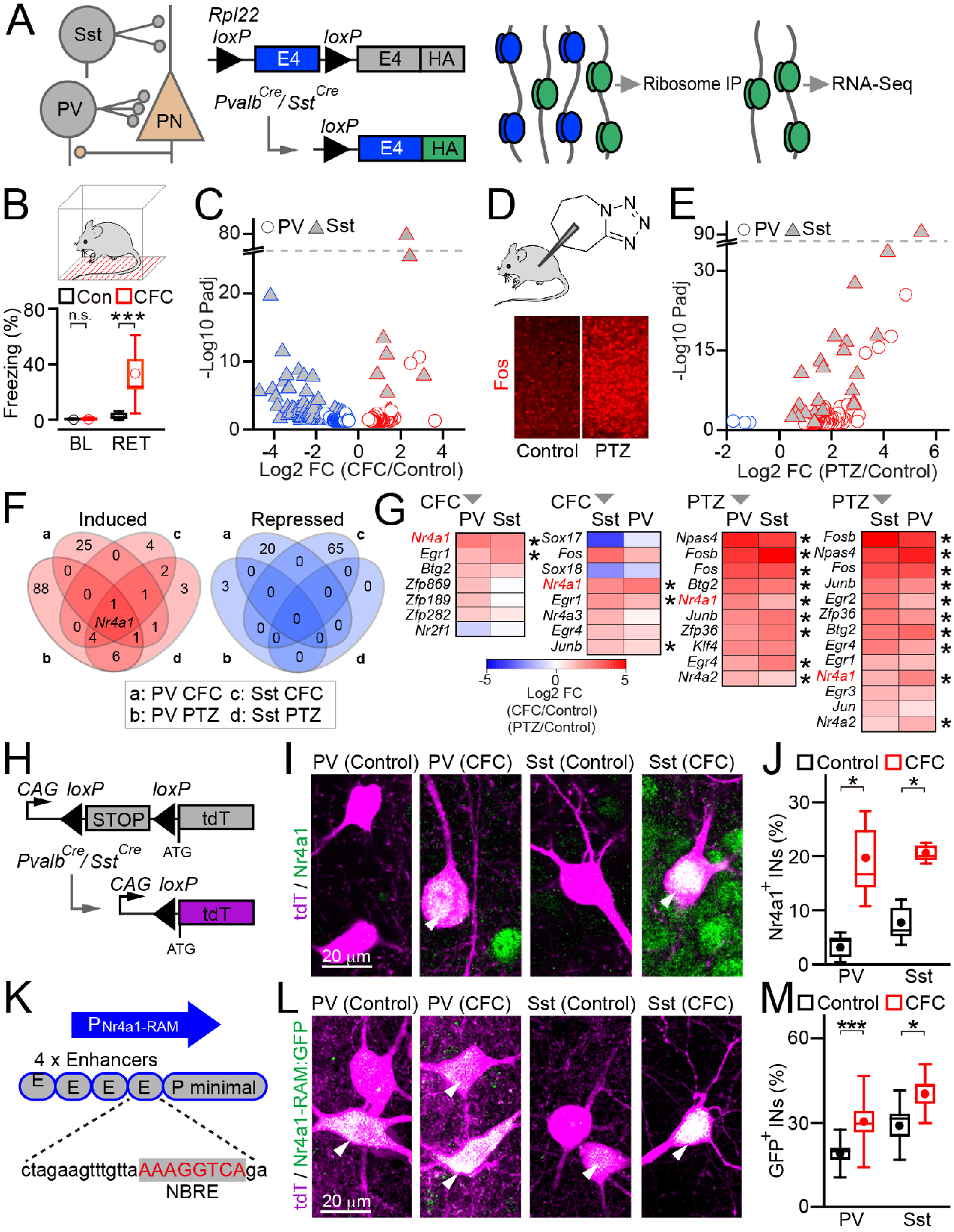
Nr4a1 is induced in PV^+^ and Sst^+^ INs during learning. **(A** to **G)** RNA-seq analysis of activity-dependent transcription. **(A)** Schematics of IN wiring with PNs and experimental workflow for mRNA profiling with genetically targeted RiboTag. **(B)** Memory retrieval in fear conditioned mice, measured as percentage of freezing under the baseline settings (BL) or in the same context (RET). *n* = 7 to 8 mice per group. **(C)** Volcano plot of differentially expressed mRNAs in PV^+^ and Sst^+^ cells 30 minutes after CFC (FC = fold change). *n* = 3 p60 mice/group. **(D)** Pharmacological induction of network activity with PTZ (i.p. injection, 50 µg/gm). Confocal images show Fos immunofluorescence in cortical columns. **(E)** Volcano plot of differentially expressed mRNAs 30 minutes after PTZ injection. *n* = 3 to 4 p60 mice/group. **(F)** Venn diagrams of significantly up- and down-regulated genes for indicated experimental settings. **(G)** Heat maps of mRNA levels of TFs. Pairwise comparisons are ranked by fold change in each IN population (arrows). Asterisks mark TFs whose expression was significantly altered in both populations. **(H)** Cre-dependent labeling of PV^+^ and Sst^+^ cells with the *Ai9* tdTomato reporter. **(I** and **J)** Confocal images of Nr4a1 immunofluorescence and quantifications of Nr4a1-positive INs in the CA1 of home-caged (Control) and fear-conditioned *Pvalb*^*Cre*^*/Ai9* and *Sst*^*Cre*^*/Ai9* reporter mice (30 minutes post-CFC). *n* = 3 mice/group. **(K)** Schematics of Nr4a1-RAM reporter. **(L** and **M)** TF activity of native Nr4a1 was monitored in CA1 INs with genetically targeted Nr4a1-RAM:DIO-GFP. Typical images and quantifications of GFP-positive INs in home-caged (Control) and fear conditioned (24 hours post-CFC) *Pvalb*^*Cre*^*/Ai9* and *Sst*^*Cre*^*/Ai9* mice are shown. *n* = 3 mice/group. In panels B, J and M, graphs represent mean values (circles), standard errors (boxes), standard deviations (whiskers), and medians (horizontal lines). * *p* <0.05; *** *p* <0.01 (defined by t-test).

To elucidate the physiological role of Nr4a1 in inhibitory circuits, we focused on hippocampal area CA1. In this brain region, PV^+^ and Sst^+^ INs participate in acquisition and retrieval of episodic memories, and their presynaptic terminals are clearly segregated in separate layers with PN somas and dendritic fields (*8, 20-22*). To confirm that Nr4a1 mRNA is translated, we conducted immunofluorescent imaging of brain sections from p60 *Pvalb*^*Cre*^*/Ai9* and *Sst*^*Cre*^*/Ai9* reporter mice in which INs were permanently labeled with tdTomato (Fig. 1H). In agreement with results of RNA-seq, CFC acutely (within 30 minutes) increased the fractions of Nr4a1-immunoreactive PV^+^ and Sst^+^ cells by 6.3- and 2.6-fold, respectively (Fig. 1, I and J). To test whether native Nr4a1 is capable of driving transcription, we designed a fluorescent reporter, Nr4a1-RAM:GFP, adapting the recently described strategy for monitoring the TF activities of Fos and Npas4 (*23*). Nr4a1-RAM was comprised of 4 short Nr4a1 enhancer repeats followed by a minimal promoter and reliably sensed pharmacologically augmented synaptic excitation or presence of the exogenous Nr4a1 in primary neuronal cultures (Fig. 1K and fig. S3, A and B). We introduced Nr4a1-RAM:GFP into PV^+^ and Sst^+^ cells of adult *Pvalb*^*Cre*^*/Ai9* and *Sst*^*Cre*^*/Ai9* mice *in vivo* with a Cre-dependent adeno-associated virus (AAV) and then imaged labeled INs in brain slices 7 days later (fig. S3C). Again, the fractions of GFP^+^ INs were significantly increased in the CA1 after CFC (24 hours post-training), albeit in this case the effect was not as pronounced as acute CFC-dependent synthesis of Nr4a1 mRNA and protein (1.6- and 1.4-fold) since 19% to 28% of cells had detectable GFP fluorescence in untrained animals, most likely because of history of their prior experience (Fig. 1, L and M and fig. S3D). Taken together, these results suggest that transient induction of Nr4a1 by external sensory cues mediates a sustained transcriptional response in INs.

Next, we generated two conditional knockout (cKO) mouse lines that lacked Nr4a1 in PV^+^ and Sst^+^ cells (*Pvalb*^*Cre*^*/Nr4a1* ^*loxP/loxP*^ and *Sst*^*Cre*^*/Nr4a1* ^*loxP/loxP*^; fig. S4). The homozygous offspring of both cKO lines were born at the expected Mendelian ratio, had regular lifespans, and did not display apparent behavioral abnormalities in the standard laboratory environment. However, quantifications of contextual freezing 2 and 24 hours after CFC demonstrated that Nr4a1-deficient mutants had aberrant short- and long-term associative memory (STM and LTM). Compared to their p60 Nr4a1 wildtype (WT) littermates, *Pvalb*^*Cre*^*/Nr4a1*^*loxP/loxP*^ cKOs exhibited prolonged freezing in the same context, though this effect was only statistically significant for STM. In contrast, the freezing times of *Sst*^*Cre*^*/Nr4a1*^*loxP/loxP*^ mice were significantly reduced at both timepoints, suggesting that STM and LTM were partially impaired (Fig. 2A). These phenotypes were not attributed to abnormal locomotion or elevated anxiety, as evidenced by measurements of travel distances and zone preference of cKOs in open fields (Fig. 2B). Furthermore, confocal imaging of brain sections from adult cKOs carrying the *Ai9* tdTomato reporter allele showed no detectable changes in the densities, positioning and gross morphologies of labeled PV^+^ and Sst^+^ cells across the hippocampal subfields, indicating that their migration, differentiation and survival were preserved (fig. S5).

**Fig. 2.**
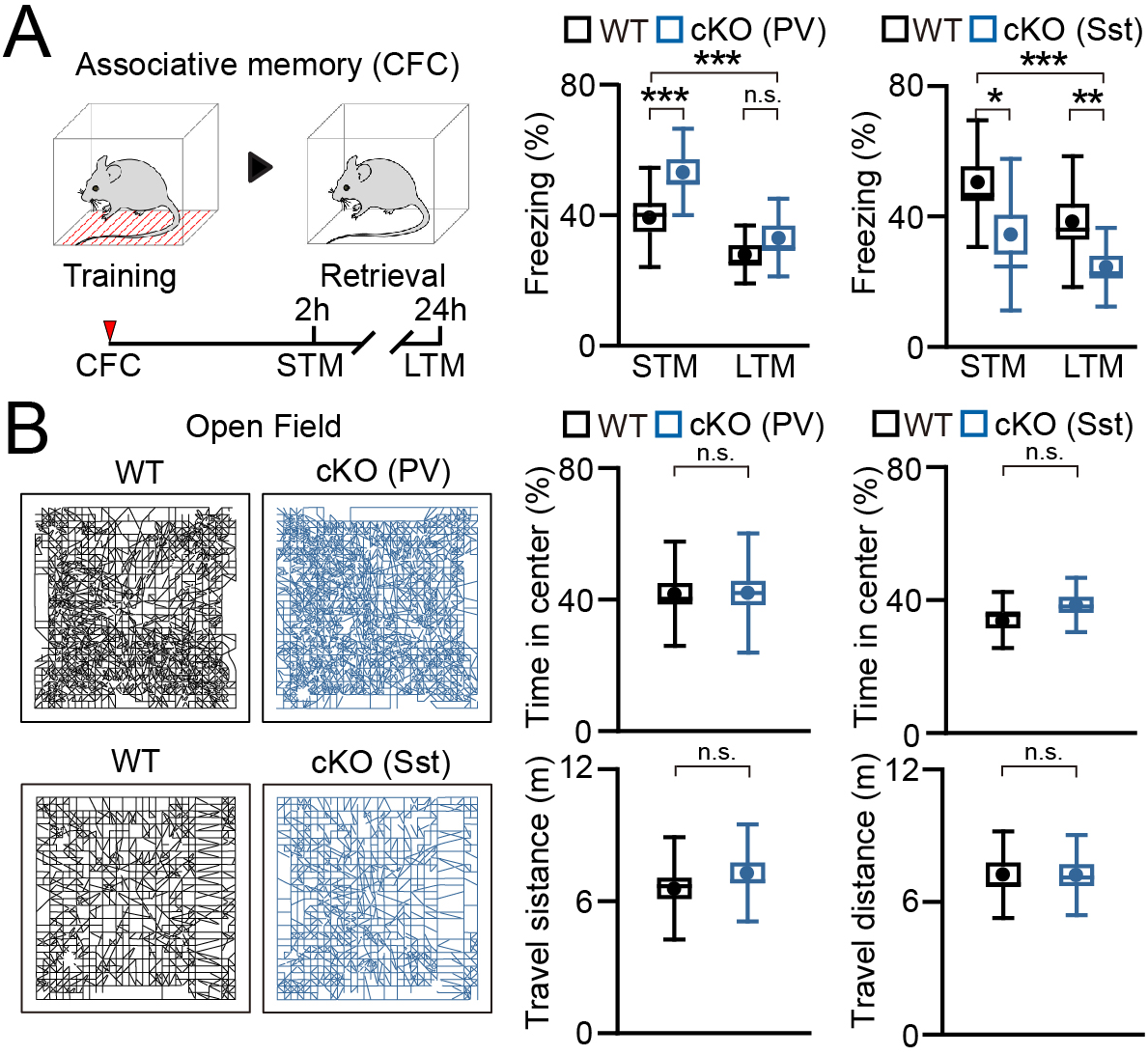
Behavior of PV^+^ and Sst^+^ Nr4a1 conditional knockout (cKO) mice. **(A)** Analysis of associative fear memory. Panel shows the experimental workflow and quantifications of retrieval of short- and long-term memory (STM and LTM), expressed as percentages of freezing in the same context 2 and 24 hours, respectively, after CFC. PV^+^(WT), *n* = 16 mice; PV^+^ (cKO), *n* = 16; Sst^+^ (WT), *n* = 17; Sst^+^ (cKO), *n* = 18. **(B)** Behavior in open fields. Panel shows typical tracks of individual animals during 30 minute recording sessions, quantifications of time spend in central zones, and total travel distances. PV^+^ (WT), *n* = 35; PV^+^ (cKO), *n* = 34; Sst^+^ (WT), *n* = 19; Sst^+^ (cKO), *n* = 18. All graphs represent mean values (circles), standard errors (boxes), standard deviations (whiskers), and medians (horizontal lines). * *p* < 0.05; ** *p* < 0.02; *** *p* <0.01 (defined by t-test and ANOVA).

To explore the possibility that Nr4a1 signaling in INs is important for GABAergic inhibition of principal projection neurons, we injected *Pvalb*^*Cre*^*/Nr4a1*^*loxP/loxP*^ and *Sst*^*Cre*^*/Nr4a1*^*loxP/loxP*^ cKOs with a Cre-dependent AAV encoding Channelrhodopsin2 (ChR2) and then sampled evoked and spontaneous inhibitory postsynaptic currents (IPSCs) from PNs in acute hippocampal slices at p21 (Fig. 3A). Whole-cell recordings of synchronous IPSCs elicited by optical stimulation demonstrated that inhibitory synaptic input to PNs from Nr4a1-deficient PV^+^ cells was significantly weaker, whereas the input from Nr4a1-deficient Sst^+^ cells was stronger compared to INs from Nr4a1 WT littermates (Fig. 3B). Likewise, PNs of *Pvalb*^*Cre*^*/Nr4a1*^*loxP/loxP*^ and *Sst*^*Cre*^*/Nr4a1*^*loxP/loxP*^ cKOs had reduced and increased frequencies of spontaneous miniature IPSCs, respectively (Fig. 3C). Nonetheless, the kinetics and amplitudes of quantal GABA currents were unaffected (fig. S6, A and B), indirectly suggesting that observed changes in inhibitory synaptic strength reflect aberrant innervation of PNs. To test this definitively, we labeled the axonal terminals of INs *in vivo* via viral Cre-dependent expression of the presynaptic marker, Synaptophysin (SyP)-Venus, and inspected their distribution at p60 across the CA1 stratum oriens (so) to pyramidal cell layer (sp) to stratum radiatum (sr) axis by confocal imaging of brain sections. Axons of INs in two mutant mouse lines projected to correct zones but their presynaptic networks were altered in opposite directions, mirroring the outcomes of electrophysiological recordings. Indeed, Nr4a1-deficient PV^+^ cells formed ∼2 times less terminals on PN somas in the CA1sp, while the innervation of PN dendritic fields by Nr4a1-deficient Sst^+^ cells was more robust. We confirmed these results by immunostaining of brain sections with antibodies against the native presynaptic marker of PV^+^ basket cells, Syt2, and pan-inhibitory presynaptic marker, GAD67 (Fig. 3, E and F; figs. S7 and S8). The presynaptic abnormalities of mutant mice were not associated with global delay in IN development or overgrowth since Nr4a1 cKO did not alter the frequencies of spontaneous glutamatergic excitation of INs or morphologies of dendritic trees of PV^+^ cells, as visualized with membrane-bound GFP (fig. S6, C to E and fig. S9). We were unable to quantify dendritic arborization of Sst^+^ cells because their dendrites intermingled with axons. To extend these studies using a complementary approach, we selectively overexpressed the full-length Nr4a1 cDNA in PV^+^ and Sst^+^ cells on the WT background from a Cre-dependent AAV and found that cell-autonomous Nr4a1 gain-of-function in INs altered the innervation of PN somas and dendrites in directions that were opposite to phenotypes cKO mice (fig. S10). Hence, our experiments with loss- and gain-of-function mouse models support the hypothesis that Nr4a1 is involved in transcriptional control of inhibitory circuit structure.

**Fig. 3.**
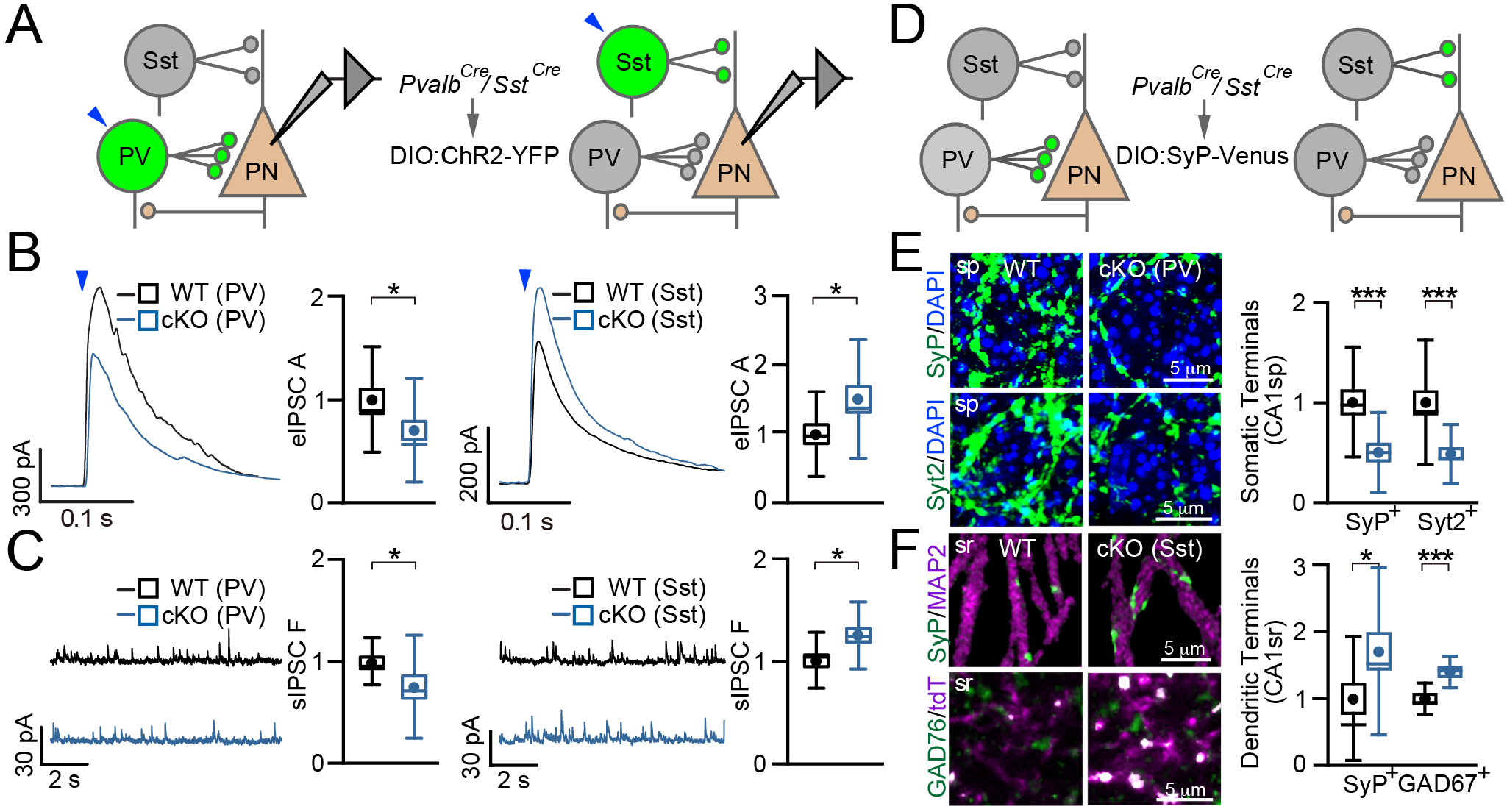
Nr4a1 exserts opposite effects on GABAergic outputs from PV^+^ and Sst^+^ INs to PNs. **(A** to **D)** Inhibitory postsynaptic currents (IPSCs) were monitored from CA1 PNs of Nr4a1 WT and IN-specific cKO mice in acute hippocampal slices. **(A)** Schematics of Cre-dependent expression of ChR2-YFP and recording configurations. **(B)** Sample traces of IPSCs evoked by optical stimulation (1 ms, 450 nm) and quantifications of eIPSC amplitudes. PV^+^(WT), *n* = 5 mice/24 neurons ; PV^+^ (cKO), *n* = 5/32; Sst^+^ (WT), *n* = 3/20; Sst^+^ (cKO), *n* = 3/21. **(C)** Sample traces of spontaneous IPSCs and quantifications of event frequencies. PV^+^ (WT), *n* = 6 mice/25 neurons; PV^+^ (cKO), *n* = 6/21; Sst^+^ (WT), *n* = 3/22; Sst^+^ (cKO), *n* = 3/25. **(D** to **F)** Aberrant innervation of PNs by Nr4a1-defieicnt INs. **(D)** Schematics of Cre-dependent labeling of presynaptic terminals with SyP-Venus. **(E)** Ablation of Nr4a1 in PV^+^ cells diminishes their innervation of PN somas. Panels show confocal images of terminals labeled in the CA1sp of Nr4a1 WT and cKO mice with SyP-Venus or antibody against Syt2 and quantifications of terminal numbers. **(F)** Ablation of Nr4a1 in Sst^+^ cells promotes their innervation of PN dendrites. Panels show confocal images of terminals labeled in the CA1sr of Nr4a1 WT and cKO mice with SyP-Venus or antibody against GAD67 and quantifications of terminal numbers. In the top row, dendrites were stained with MAP2. In the bottom row, IN projections were labeled with the *Ai9* tdTomato reporter. (*n* = 3 to 4 mice/genotype). All graphs represent mean values (circles), standard errors (boxes), standard deviations (whiskers), and medians (horizontal lines). * *p* < 0.05; *** *p* <0.01 (defined by t-test). cKO values were normalized to WT for each littermate pair.

Neuronal connectivity is organized by secreted and transmembrane proteins that guide axons to receptive fields and mediate surface adhesion or repulsion (*1*). These proteins are often expressed in a cell-type-specific manner, thereby generating combinatorial codes that define the wiring selectivity of diverse neuron classes and/or their synapse numbers in local connectomes (*2, 3, 24*). To determine how Nr4a1 regulates the presynaptic networks of INs at a molecular level, we performed *in vivo* screens for downstream effectors of this TF in PV^+^ and Sst^+^ cells. To this end, we generated cKO mouse lines carrying the RiboTag allele (*Pvalb*^*Cre*^*/Nr4a1*^*loxP/loxP*^*/RiboTag* and *Sst*^*Cre*^*/Nr4a1*^*loxP/loxP*^*/RiboTag*) and surveyed transcriptional profiles of Nr4a1-deficient INs at p60 by RNA-seq. Analysis with the cutoff criteria of absolute FC of expression level >2 yielded 171 and 269 putative Nr4a1 targets in PV^+^ and Sst^+^ cells, respectively. Approximately halves of these genes were upregulated in mutant mice, indicating that Nr4a1 may act on different substrates as a transcriptional activator and repressor (Fig. 4A). Remarkably, ablation of Nr4a1 produced a partial switch in lineage-specific genetic programs by inducing genes that are repressed and, *vice versa*, by repressing genes that are transcribed in a particular IN lineage in the normal brain (Fig. 4B). For example, of 83 genes whose mRNA levels were >2-fold lower in Nr4a1-deficient PV^+^ cells, nearly 80% had enriched mRNAs in their WT counterparts comparative to WT Sst^+^ cells. Consistent with these findings, >96% of all Nr4a1 effectors were lineage-specific (Fig. 4C). Bioinformatic searches showed that many of these genes contained Nr4a1 consensus DNA binding sites (*25*) and encoded proteins with axonal and synaptic functions (fig. S11).

**Fig. 4.**
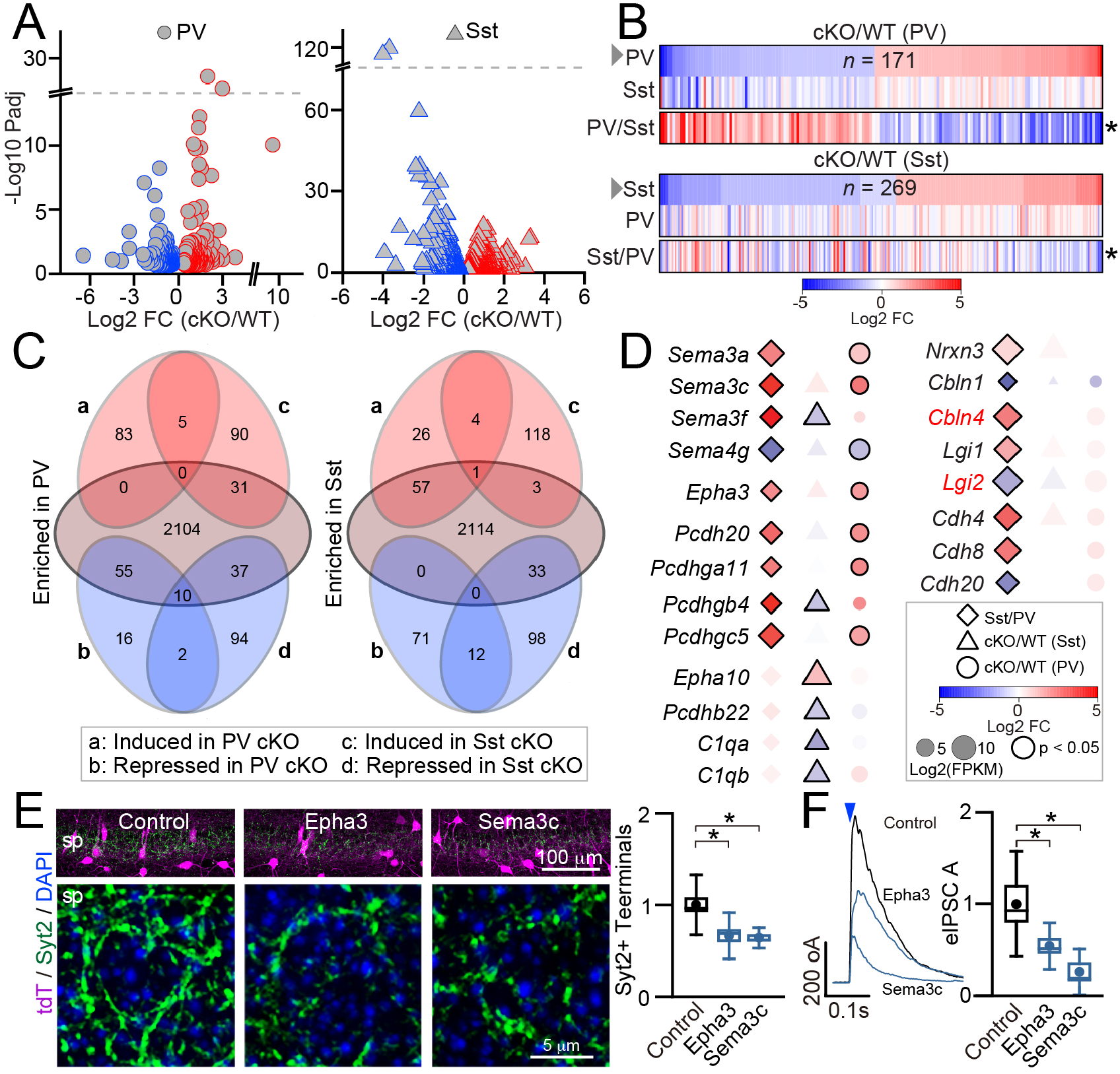
Nr4a1 regulates cell-specific molecular programs essential for IN wiring. **(A** to **D)** Transcriptional profiles of PV^+^ and Sst^+^ INs, as assessed by deep sequencing of mRNAs isolated from Nr4a1 WT and cKO mice with genetically targeted RiboTag (*n* = 3 mice/group). **(A)** Volcano plots of RNA-seq data (FC = fold change). **(B)** Heat maps show comparisons of mRNA levels of genes that were significantly up- or down-regulated in each population of Nr4a1-deficient INs (top rows, FDR<0.05, -1<log2 FC>1), with their expression in other Nr4a1-deficient population (middle rows) and relative expression in Nr4a1 WT mice (bottom rows). **(C)** Venn diagrams for data shown in panel (B). **(D)** Loss of Nr4a1 leads to a switch in cell-specific expression of genes involved in synaptic differentiation. Examples of affected (left) and unaffected (left) genes from families encoding surface adhesion/repulsion molecules are shown. Annotated heatmaps demonstrate relative expression levels in normal INs (first columns, plotted as Sst/PV ratios) and expression in each population of Nr4a1-deficient INs (second and third columns). **(E** and **F)** Gain-of-function of developmentally repressed *Epha3* and *Sema3c* mimics the effects of Nr4a1 cKO in PV^+^ INs. **(E)** *Epha3* or *Sema3c* cDNAs were expressed in the CA1 of *Pvalb*^*Cre*^*/Ai9* mice from Cre-dependent AAVs. Panels shows confocal images of brain sections labeled with the antibody against Syt2 and quantifications of Syt2-positive terminals in CA1sp. Control, *n* = 3 mice/24 sections; Epha3, *n* = 3/20; Sema3c, *n* = 3/21. **(F)** *Epha3* or *Sema3c* were co-expressed in PV^+^ cells with ChR2-YFP. Panels shown sample traces of light-induced eIPSCs recorded from PNs and quantifications of eIPSC amplitides. Control, *n* = 3 mice/9 neurons; Epha3, *n* = 3/13; Sema3c, *n* = 3/9.

To address the possibility that consequences of Nr4a1 signaling on IN connectivity with PNs reflect the role of Nr4a1 in controlling the synaptic organizing code, we next compared the mRNA levels of genes from families essential for axonal and/or synaptic differentiation in p60 WT and cKO mice. Extending recent studies in the developing cerebral cortex (*2*), we found that PV^+^ and Sst^+^ cells in the adult brain expressed different combinations of Neurexins (*Nrxn*), Neuroligins (*Nlgn*), Cerebelins (*Cbln*), Leucine-rich glioma-inactivated (*Lgi*), classical Cadherins (*Cdh*), Protocadherins (*Pcdh*), Semaphorins (*Sema*), Ephrin receptors (*Epha*), Complement 1q (*C1q*), Contactins (*Cntn*), Contactin associated proteins (*Cntnap*), Fibroblast growth factors (*Fgf*), Immunoglobulin superfamily proteins (*Igsf*), Neurexophilins (*Nxph*) and Latrophilins (*Adgrl*). These unique patterns were disrupted in the absence of Nr4a1 due to pronounced switches in lineage-specific transcription of *Sema, Epha, Pcdh* and *C1q* while other genes were either unaffected or affected to a lesser extent. Notably, Nr4a1-deficient INs had preserved combinatorial expression of *Cbln* and *Lgi* whose members influence the subcellular specificity of synaptic targeting (*2*) (Fig. 4D and fig. S12,A and B).

Transcriptional switches detected in Nr4a1 mutant mouse lines by RNA-seq are reconcilable with effects of Nr4a1 loss- and gain-of-function on IN wiring since the members of *Sema, Epha, Pcdh* and *C1q* gene families can mediate attraction and repulsion in various contexts (*1, 26-28*). To further establish a mechanistic link between Nr4a1 and its effectors, we focused on *Sema3c* and *Epha3*. Both of these genes had enriched mRNAs in WT Sst^+^ cells, both contained Nr4a1 consensus DNA binding motifs in promoter regions upstream transcription start sites, and both were strongly and selectively de-repressed in PV^+^ cells lacking Nr4a1 (Fig. 4D and fig. S12C). We therefore predicted that Nr4a1 restricts the somatic innervation of PNs by PV^+^ basket cells by restricting the transcription of *Sema3c* and *Epha3* which act as local repulsive cues. To test this model experimentally, we introduced recombinant Sema3c and Epha3 cDNAs into CA1 PV^+^ cells of *Pvalb*^*Cre*^*/Ai9* reporter mice *in vivo* with Cre-dependent AAVs on the Nr4a1 WT background. Confocal imaging of tdTomato and Syt2-immunoreactive presynaptic terminals showed no changes in the survival of INs overexpressing individual constructs, their distribution, and axonal targeting to CA1sp. However, the numbers of inhibitory synapses on PN somas were reduced in both cases, as revealed by quantitative measurements of Syt2^+^ puncta density and whole-cell recordings of synchronous IPSCs elicited in PNs by optogenetic stimulation of PV^+^ cells (Fig 4. E and F). Thus, *Sema3c* and *Epha3* gains-of-function mimic the phenotypes of Nr4a1 cKO.

Herein, we investigated transcriptional programs that regulate synaptic inhibition in the mouse brain. Our results demonstrate that the nuclear receptor encoded by an early response gene, *Nr4a1*, is induced in GABAergic INs during associative learning and is required for normal memory storage. At the cellular level, Nr4a1 can promote or suppress the local axonal connectivity of INs that provide GABAergic inputs onto distinct subcellular compartments of their postsynaptic partners. At the molecular level, Nr4a1 instructs the axonal wiring, at least in part, through repertoires of cell-specific effectors mediating surface adhesion and repulsion. A large body of work shows that assembly of synaptic contacts in the developing brain is coordinated by numerous proteins localized in axons and dendrites, and that knockouts of individual players in whole animals often result in only minor phenotypes (*1*). For example, the bulk of synaptogenesis is preserved in mice lacking all isoforms of Neurexins (*29*), even though asymmetric Neurexin-Neuroligin interactions are sufficient for the formation of artificial synapses in simple heterologous systems (*30*). It is therefore plausible that molecular events underlying structural plasticity are equally complex. In agreement with recent studies of wiring specificity of excitatory and inhibitory neurons (*2, 3*), we find that INs in the adult forebrain employ non-overlapping sets of synaptic organizers. The Nr4a1 signaling resembles partial reprogramming of this code attained via on- and off-switches in several gene families. Hence, the unique combinatorial expression patterns of genes that define synaptic architecture of a given neuron are not solely determined during development and maintained across the lifespan in an intrinsic manner. Rather, these patterns can be dynamically re-arranged by inducible TFs whose transcription is regulated by sensory cues. Since ERGs are induced in activated neuronal ensembles throughout the brain (*4*), similar transcriptional switches may contribute to structural plasticity of other circuits and cell types. It is noteworthy that Nr4a1 does not exclusively control “synaptic” genes. Some of its effectors appear to act in metabolic pathways, and their roles in GABAergic INs need to be investigated in the future. As disfunctions of inhibitory circuits are associated with a broad spectrum of neurological disorders, our analyses of Nr4a1 signaling in INs may also have translational implications.

## ACKNOWLEDGEMENTS

We thank Dr. M. Mayford, Dr. G. Lippi and all members of the Maximov lab for advice and critical comments; Dr. K. Spencer for assistance with microscopy; Dr. A. Roberts for assistance with behavioral experiments; and The Scripps Research Institute (TSRI) Next Generation Sequencing Core for deep sequencing and bioinformatics support.

This study was supported by the US National Institutes of Health grants R01MH118442, R01NS087026, R01GM117049 and RF1 MH123224 to A.M.

## Author contributions

A.M. and M.H. conceived the study. M.H. generated mutant mice and carried out molecular profiling, imaging, whole-cell recordings, and behavioral experiments. S.P. performed initial RNA-seq screens with PTZ-injected mice and generated AAV expression vectors. F.P. and V. R. assessed the expression of Nr4a1 in reporter mice by immunofluorescent imaging. J.C. and G.S. assisted M.H. with genetic experiments and data analysis. A.M. wrote the manuscript.

## Data and materials availability

RNA seq data will be deposited to GEO. All reagents, including unique reporters and expression vectors can be obtained from A.M. upon request.

**SUPPLEMENTARY MATERIALS** are enclosed with the manuscript and include detailed description of materials and methods, 12 figures and figure legends.

## Supplementary Materials

### Materials and Methods

#### Mice

The *Pvalb*^*Cre*^, *Sst*^*Cre*^, *Nr4a1*^*loxP/loxP*^, *RiboTag* and *Ai9* mouse alleles used in this study have been described previously (*10-12, 31, 32*). Mice were housed, crossed, and analyzed according to protocols approved by the IACUC committee. All animals were a mix of C57BL/6 and 129/SV backgrounds. Males and females were examined together, with exception of behavior.

### Expression vectors

Adeno-associated viruses (AAVs) were produced with well-characterized shuttle vectors containing the EF1α promoter and the DIO cassette for inducibility with Cre (*33-35*). For targeted overexpression of Epha3 and Sema3C cDNAs, the 1.26 kb EF1α promoter was replaced with the 212 bp EF1α-core promoter to maintain the overall sizes of inserts within the packaging limit (∼4.4kb). This smaller promoter was amplified by PCR from the pJEP308 plasmid (Addgene #113685). The coding sequences of full length Epha3, Sema3C and Nr4a1 were amplified by PCR from the custom-made mouse hippocampal cDNA library with the following primers:

<u>Epha3</u>

Forward: 5′-GCTAGCATGGATTGTCACCTCTCCATCCT-3′

Reverse: 5′-TTAATTAATTACACTGGAACTGGACCATTCTTAGATTG-3′

<u>Sema3C</u>

Forward: 5′-GCTAGCATGGCATTCCGGACAATTTGC-3′

Reverse: 5′-TTAATTAATTATGACTCTGGCAACTGATTCCTCCT-3′

<u>Nr4a1</u>

Forward: 5′-CGGATCCATGCCCTGTATTCAAGC-3′ Reverse: 5′-GGAATTCGAAAGACAATGTGTCCAT-3′

The Epha3 and Sema3C coding sequences were inserted downstream of EF1α-core promoter in the 3′– 5′ orientation using PacI and NheI restriction sites and were flanked by two pairs of loxP sites (DIO). The Nr4a1 coding sequence was inserted in the same manner into AAV-EF1α-DIO-GFP vector using BamHI and EcoRI sites.

The Nr4a1-RAM reporter was constructed according to previously described strategy for reporters of Fos and Npas4 (*23*). This reporter contained 4 tandem Nr4a1 enhancer repeats of the following sequence with underlined nucleotides representing the binding motif:

5′-CTAGAAGTTTGTTA<u>AAAGGTCA</u>GA -3′

The Fos minimal promoter was amplified by PCR from the pAAV-CRAM-tdTomato plasmid (Addgene #84468). The enhancer modules were inserted upstream of this promoter and the entire cassette was then used to replace the EF1α promoter in the EF1α:DIO-GFP AAV shuttle vector.

The AAV/LV shuttle vectors for expression of mGFP, SyP-Venus, ChR2-EYFP, Cre and ΔCre have been described previously (*33, 36, 37*).

### AAV production and injection

AAVs were generated in house in HEK293T cells, purified by Heparin-based affinity chromatography, tittered by real-time quantitative PCR, and injected into brains of mice carrying the Cre drivers at titers of 2×10^12^ GC/ml as we have previously described (*33-35*). For synaptic tracing, reconstruction of neuronal morphologies, optogenetic stimulation and overexpression of cDNAs, AAVs were delivered to neonates. Pups were anesthetized on ice, bilaterally injected into the cerebral lateral ventricles with 1 µl of viral stocks using a glass micropipette (1 mm O.D., 0.5 mm I.D., 10 µm tip diameter), warmed up for 5 minutes on a heat pad, and returned to home cages until experiments. To monitor the transcriptional activity of Nr4a1 with Nr4a1-RAM:GFP, the reporter virus was delivered to young adults and imaged 7 days later. Mice were anesthetized with 1.5%–2% isoflurane in O_2_; stereotaxic injections were performed bilaterally into the dorsal CA1 with the following coordinates (relative to bregma) and volumes: AP −1.82 mm, ML ± 1.4 mm, DV −1.2 mm. Viruses were infused for 5 minutes at a rate of 100 nl per minute.

### LV production and infection

Recombinant lentiviruses (LVs) were produced by co-transfection of HEK293T cells with corresponding shuttle vectors, and pVSVg, pGag-Pol and pRev plasmids that encode the elements essential for packaging of viral particles. Transfections were performed using FuGENE HD reagent (Roche). Conditioned medium with secreted viruses was collected 48 hours later and centrifuged at 5,000g for 5 minutes to remove cellular debris. Neurons were infected with 200 μl of viral supernatant per each well of a 24-well plate at 4 days *in vitro* (DIV). This protocol enabled reliable infection of ∼95% neurons in culture.

### Neuronal cultures

Cortices of p0 pups were dissociated with papain and seeded onto 24-well plates coated with poly-D-Lysine (Millipore). Cultures were maintained for 1 hour in MEM (Invitrogen) supplemented with fetal bovine serum (Invitrogen), and glucose (Sigma), followed by incubation in the serum-free Neurobasal-A (Invitrogen) supplemented with B27 (Invitrogen) and Glutamax (Invitrogen). Cultures were kept at 37°C in humidified incubators with 5% CO2 until use.

### Validation of *Nr4a1*^*loxP/loxP*^ allele and Nr4a1-RAM reporter *in vitro*

Neurons from *Nr4a1*^*loxP/loxP*^ neonates were infected at DIV4 with LVs expressing Cre or catalytically inactive ΔCre under the control of the Ubiquitin (Ubc) promoter and assayed 3 days later. Genotyping was performed by PCR with the following primers:

Forward: 5′-TGACACCCTCACACGGACAA-3′

Reverse: 5′-CCAGTACATAGAGGATGCTTGTT-3′

Cre-dependent ablation of Nr4a1 was also confirmed by qPCR of reverse transcribed mRNAs with the following primers:

<u>Fos</u>

Forward: 5′-GGGACAGCCTTTCCTACTACC-3′

Reverse: 5′-AGATCTGCGCAAAAGTCCTG-3′

<u>Nr4a1</u>

Forward: 5′-TTGAGTTCGGCAAGCCTACC-3′

Reverse: 5′-GTGTACCCGTCCATGAAGGTG-3′

<u>Gapdh</u>

Forward: 5′-TCAACGGGAAGCCCATCA-3′

Reverse: 5′-CTCGTGGTTCACACCCATCA-3′

To validate the Nr4a1-RAM reporter, neurons from C57BL/6 neonates were infected at 4DIV with LV Ubc-Cre, AAVDJ Nr4a1-RAM:DIO-GFP and AAVDJ EF1α:DIO-Nr4a1. At 14DIV, cultures were stimulated overnight with Bicuculline (50 μM, Sigma) and 4-aminopyridine (250 μM, Tocris). Cultures were then rinsed briefly with HBSS (Invitrogen) and reporter expression was assessed by fluorescent imaging.

### Drug injection

Pentylenetetrazol (PTZ, Sigma) was dissolved in saline prior to each experiment and administered to mice by intraperitoneal injections through a 29 g needle at a dose of 50 mg/kg body weight. Control vehicle solution contained saline alone.

### mRNA isolation with RiboTag

Cortices of p60 *Pvalb*^*Cre*^/*RiboTag* and *Sst*^*Cre*^/*RiboTag* mice were dissected in ice-cold PBS and homogenized in the buffer containing 100 mM KCl, 50 mM Tris-HCl pH 7.4, 12 mM MgCl2, 100 µg/ml cycloheximide (Sigma), 1 mg/ml heparin (Sigma), 1x complete mini, EDTA-free protease inhibitor cocktail (Roche), 200 units/ml RNasin© plus inhibitor (Promega) and 1 mM DTT (Sigma) (500 µl ∼3% w/v). The lysate was centrifuged at 2,000g for 10 minutes at 4°C. Igepal-CA380 was then added to the supernatant at the final concentration of 1%. The lysate was briefly incubated on ice and then centrifuged at 12,000g for 10 minutes at 4°C. 25 µl of the supernatant was collected as input sample. 30 µl/ml of anti-HA coupled magnetic beads (Pierce) were then added to the remaining supernatant. Incubation was performed on a rotator for 3 hours at 4°C. The beads were washed four times in the high salt buffer containing 350 mM KCl, 1% Igepal-CA380, 50 mM Tris-HCl, pH7.4, 12 mM MgCl2, 100 µg/ml Cycloheximide (Sigma) and 1 mM DTT (Sigma). Ribosomes were eluted in 350 µl of RLT plus buffer (Qiagen). RNA purification was performed using RNeasy Plus Mini kit (Qiagen) following the manufacturers’ instructions. For qPCR, 20 ng of total RNA was reverse transcribed using High-Capacity cDNA Reverse Transcription Kit (Applied Biosystems). qPCR quality controls were performed with the following primers:

<u>Fos</u>

Forward: 5′-GGGACAGCCTTTCCTACTACC-3′

Reverse: 5′-AGATCTGCGCAAAAGTCCTG-3′

<u>Nr4a1</u>

Forward: 5′-TTGAGTTCGGCAAGCCTACC-3′

Reverse: 5′-GTGTACCCGTCCATGAAGGTG-3′

<u>Gapdh</u>

Forward: 5′-TCAACGGGAAGCCCATCA-3′

Reverse: 5′-CTCGTGGTTCACACCCATCA-3′

<u>Syt2</u>

Forward: 5′-AGAACCTGGGCAAATTGCAGT-3′

Reverse: 5′-CCTAACTCCTGGTATGGCACC-3′

<u>Pvalb</u>

Forward: 5′-CATTGAGGAGGATGAGCTG-3′

Reverse: 5′-AGTGGAGAATTCTTCAACCC-3′

<u>Lhx6</u>

Forward: 5′-CTACTTCAGCCGATTTGGAACC-3′

Reverse: 5′-GCAAAGCACTTTCTCCTCAACG-3′

### Deep sequencing

RNA-seq was performed at the TSRI Next Generation Sequencing Core on the Illumina HiSeq platform. The libraries were generated, barcoded, and sequenced according to the manufacturers’ recommendations. The reads were trimmed from adapter sequences using cutadapt 1.18 with Python 3.6.3. The trimmed reads were mapped to the reference genome using the STAR aligner 2.5.2a and gene abundance was estimated with python 2.7.11, and HTSeq 0.11.0. Differential expression analysis was performed in the R package DESeq2. DESeq2 first adjusts read counts based on a normalization factor that accounts for sample size. This was followed by dispersion estimates based on a negative binomial model which accounts for genes with very few counts. Finally, the Wald test was performed to test for statistical significance. Genes with adjusted p-values (padj) of <0.05 were identified as differentially expressed. To remove genes with high fold changes due to low expression, a minimum normalized expression level (basemean) filter of 50 was added. The filtered data were used to generate volcano plots and heatmaps.

### Bioinformatics

Pathway analyses were performed using Gene Ontology and STRING. The STRING networks were generated with the confidence score of 0.7. Putative Nr4a1 DNA binding motifs proximal to transcription start sites (TSS) of target genes were identified in HOMER v4.11 with the function findMotifs.pl and the criteria ‘-start -2000 -end 100 -len 8,10 -p 4’. For comparative analysis of cell-specific genetic programs in normal and Nr4a1-deficient neurons, dot plots were generated using the “ggplot2” R package. In these plots, different symbols were used to mark cell populations and/or enrichment. Symbol sizes corresponded to log2 of average expression level (log2(basemean)) of each gene. Differentially expressed genes were presented with bold outlines (padj<0.05). Up- and down-regulated genes were color coded based on log2 of fold changes (log2 FC). Previously published datasets from the same IN populations at earlier developmental stages (*2*) were used as internal controls.

### Immunofluorescent and reporter imaging

For immunocytochemistry, cultured neurons attached to the glass coverslips were rinsed once in PBS, fixed for 15 minutes on ice in 4% paraformaldehyde (PFA), 4% sucrose in PBS, and permeabilized for 5 minutes at room temperature in 0.2% Triton X-100 (Roche). After permeabilization, neurons were incubated for 1 hour in the blocking solution containing 5% BSA (Sigma, fraction V) in PBS, followed by overnight incubation with primary and corresponding fluorescently labeled secondary antibodies diluted in the same blocking solution. Samples were washed three times in PBS (10 minutes per wash) after each antibody incubation. The coverslips were then mounted on glass slides with PVA-DABCO (Sigma).

For immunohistochemistry, mice were anesthetized with isoflurane and perfused transcardially with 25 ml of ice-cold PBS followed by 25 ml of 4% PFA in PBS using a peristaltic pump. The brains were removed, incubated overnight in 0.4% PFA, and sliced in ice-cold PBS using a vibratome. 90 μm thick coronal sections were briefly boiled in the citrate solution containing 10 mM sodium citrate and 0.05% Tween20 (pH 6.0) for antigen retrieval, and subsequently incubated for 5 minutes in the same solution at room temperature. Sections were then washed three times in PBS (5 minutes per wash), blocked for 3 hours in the solution containing 4% BSA, 3% donkey serum and 0.5% Triton, and incubated overnight with primary antibodies diluted in the same blocking solution. Subsequently, sections were washed three times in PBS, incubated with corresponding fluorescently labeled secondary antibodies, washed, mounted on glass slides, and embedded in PVA-DABCO (Sigma).

The following antibodies were used at indicated dilutions:

Chicken anti-GFP (1:1,000, Aves), Mouse anti-Syt2 (1:20, ZIRC), mouse anti-Nr4a1 (1:50, Santa Cruz), Rabbit anti-VGAT (1:1,000, Synaptic Systems), chicken anti-MAP2 (1:10,000, Abcam), mouse anti-GAD67 (1:1,000, Millipore) and Alexa 488, 546, 647 secondary antibodies (1:500, Invitrogen).

3D images were collected under the Nikon C2 or A1 confocal microscopes with 0.3-0.5 μm Z-intervals using 20x, 40x and 60x objectives. All images were subjected to uniform digital filtering to reduce non-specific background.

### Image data analysis

SyP-Venus or Syt2-positive presynaptic terminals of PV^+^ cells were automatically marked in Volocity software. The CA1 pyramidal cell layer was visualized by DAPI staining and manually assigned as a region of interest (ROI). Data were collected as density (number of terminals per unit of ROI volume). Threshold values were established for individual channels and applied equally across datasets. Similar strategy was used to quantify the overall numbers of terminals of Sst^+^ cells in CA1sr followed by analysis of dendritic innervation in Imaris (Bitplane). The dendrites of pyramidal cells were visualized by immunolabeling for MAP2 and then reconstructed with the “Create surface” tool to automatically calculate areas and create masks. SyP-Venus-positive boutons were detected with the “spot” tool within these masks. Data were collected as density (number of contacts per unit of dendritic surface). GAD67- and tdTomato-positive boutons were detected with the “spot” tool. Colocalization was analyzed with the “Colocalize spots” tool (Imaris XT extension). Threshold values were calculated individually for each brain using the “above automatic threshold” tool. The individual values were averaged and applied to all images. For analysis of dendrite morphologies, 3D images of single neurons labeled with mGFP were reconstructed from serial stacks and analyzed in Neurolucida (MBF Bioscience). The numbers of PV^+^ and Sst^+^ cells within different hippocampal regions of *Pvalb*^*Cre*^/*Ai9* and *Sst*^*Cre*^/*Ai9* mice were quantified manually in NIS elements (Nikon). For analysis of Nr4a1-RAM induction, cells expressing GFP and tdTomato were detected with the “Spot” tool in Imaris with a spot diameter of 18 µm and 2 additional filters (“Intensity Max Ch=tdTomato” above automatic threshold, “Intensity Median Ch=GFP” above automatic threshold). For analysis of native Nr4a1 induction, ROIs were drawn in NIS Elements (Nikon) using DAPI as a nuclear marker of PV^+^ or Sst^+^ cells labeled with tdTomato. Background ROIs were assigned in CA1sr distant from CA1sp, and their intensity values were subtracted from the values of Nr4a1 intensity. Threshold for Nr4a1-immunoreactive cells was generated based on sample background and uniformly applied to all quantifications from all samples.

### Electrophysiology

Mice were anesthetized with isoflurane. Brains were removed and placed into ice-cold oxygenated buffer (95%O2/5%CO2) containing 228 mM sucrose, 2.5 mM KCl, 0.5 mM CaCl_2_, 7 mM MgCl_2_, 26 mM NaHCO_3_, 1 mM NaH_2_PO_4_, and 11 mM glucose. Transverse, 300 μm thick slices were sectioned with a vibratome and initially stored at 32°C in oxygenated aCSF containing 119 mM NaCl, 2.5 mM KCl, 1 mM NaH_2_PO_4_, 26 mM NaHCO_3_, 1.3 mM MgCl_2_, 2.5 mM CaCl_2_, 11 mM glucose (pH 7.4, 292 mOsm), and then allowed to recover for 1 hour in oxygenated ACSF at 24 °C prior to recording. Synaptic currents were monitored in whole-cell voltage-clamp mode using Multiclamp 700B amplifier (Molecular Devices, Inc.). Recordings were performed at room temperature. The pipette solution contained 135 mM CsMeSO_4_, 8 mM CsCl, 0.25 mM EGTA, 10 mM HEPES, 2 mM MgATP, 0.3 mM Na_2_GTP, 5 mM QX-314, and 7 mM Na_2_phosphocreatine (pH 7.4, 302 mOsm). Synchronous release was triggered by 1 ms optogenetic stimulation through the Lambda DG-4 illumination system. The frequency, duration, and magnitude of stimuli was controlled by Lambda DG4 (Sutter Instrument, Inc.) and pClamp10 through TTL. Traces were analyzed offline with pClamp10 (Molecular Devices, Inc.) and OriginPro (Origin Lab) software packages.

### Behavior

#### *Open* field tests (OFT)

Locomotor activity was measured in polycarbonate cages (42 × 22 × 20 cm) placed into frames (25.5 × 47 cm) mounted with two levels of photocell beams (2 and 7 cm) above the bottom of the cage (San Diego Instruments, San Diego, CA). These two sets of beams allow for the recording of both horizontal (locomotion) and vertical (rearing) behavior. A thin layer of bedding material was applied to the bottom of the cage. Mice were tested for 120 minutes.

#### Contextual fear conditioning (CFC)

Mice were allowed to explore the fear-conditioning boxes (Context A, Med Associates SD) for 3 minutes and were then subjected to four bursts of foot shocks (0.55 mA, 1 minute inter-shock intervals). Memory tests consisted of one 3 minute exposure to the training box. Freezing was determined in 0.75 seconds bouts and expressed as percent time in the context. The following numbers of p60 control (Cre drivers with WT alleles) and cKO mice of indicated sexes were used. OFT: PV^+^ WT: *n* = 18 males and 17 females; PV^+^ cKO (*Pvalb*^*Cre*^*/Nr4a1*^*loxP/loxP*^): *n* = 16 males and 18 females; Sst^+^ WT: *n* = 6 males and 13 females; Sst^+^ cKO (*Sst*^*Cre*^*/Nr4a1*^*loxP/loxP*^): *n* = 6 males and 12 females; CFC: PV^+^ WT: *n* = 16 females; PV^+^ cKO (*Pvalb*^*Cre*^*/Nr4a1*^*loxP/loxP*^): *n* = 16 females; Sst^+^ WT: *n* = 17 males; Sst^+^ cKO (*Sst*^*Cre*^*/Nr4a1*^*loxP/loxP*^): *n* = 18 males.

### Quantifications and Statistics

All quantifications and statistical analyses were performed in OriginPro (Origin Lab). Means, medians, standard deviations (S.D.) and standard errors (S.E) are shown in all graphs. P values were determined with Student’s t-test (for two groups) and analysis of variance (for multiple groups). Welch’s correction was applied to Student’s t-test when two groups have unequal sample size. All manual quantifications were performed in a blind manner, by investigators who were unaware of genotypes and experimental conditions. Detailed descriptive statistics can be found in figure legends.

## Supplementary figures and legends

**Fig. S1.**
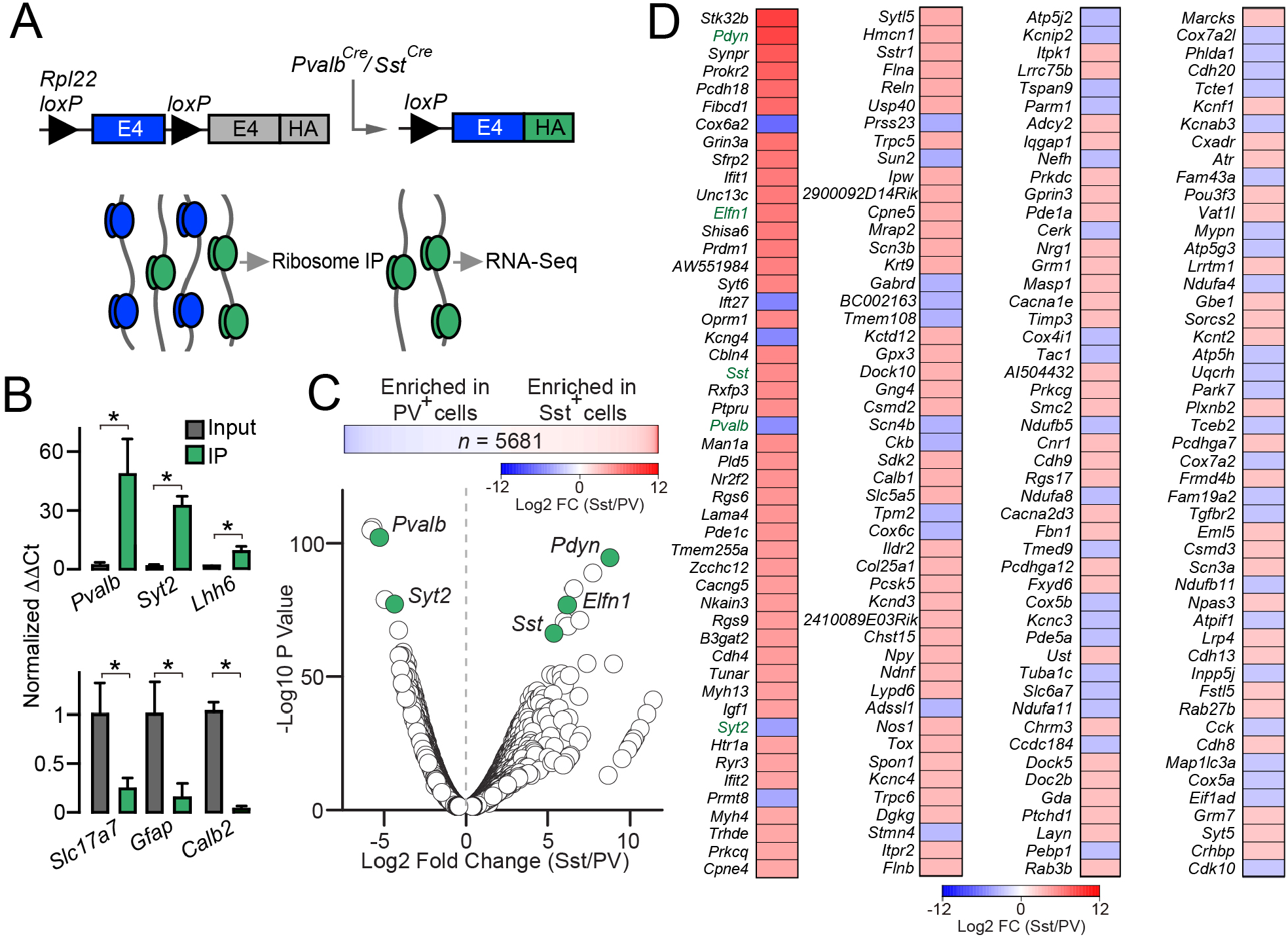
Transcriptional profiling of PV^+^ and Sst^+^ INs in p60 mice. **(A)** Experimental workflow for deep sequencing of mRNAs isolated with genetically-targeted RiboTag. **(B)** Examples of qPCR quality controls of sample purity. Graphs show normalized ΔΔCt values for markers of PV^+^ cells (top) or excitatory neurons and astrocytes (bottom) in libraries from cortices and hippocampi of *Pvalb*^*Cre*^/*RiboTag* mice (*n* = 3). **(C)** Heatmap and volcano plot of relative expression levels of 5681 genes in INs of *Pvalb*^*Cre*^/*RiboTag (n* = 2) and *Sst*^*Cre*^/*RiboTag* mice (*n* = 4) maintained in home cages. Known markers of each IN population are labeled in green. **(D)** Heatmaps of top 200 differentially expressed transcripts (plotted as Sst/PV ratios).

**Fig. S2.**
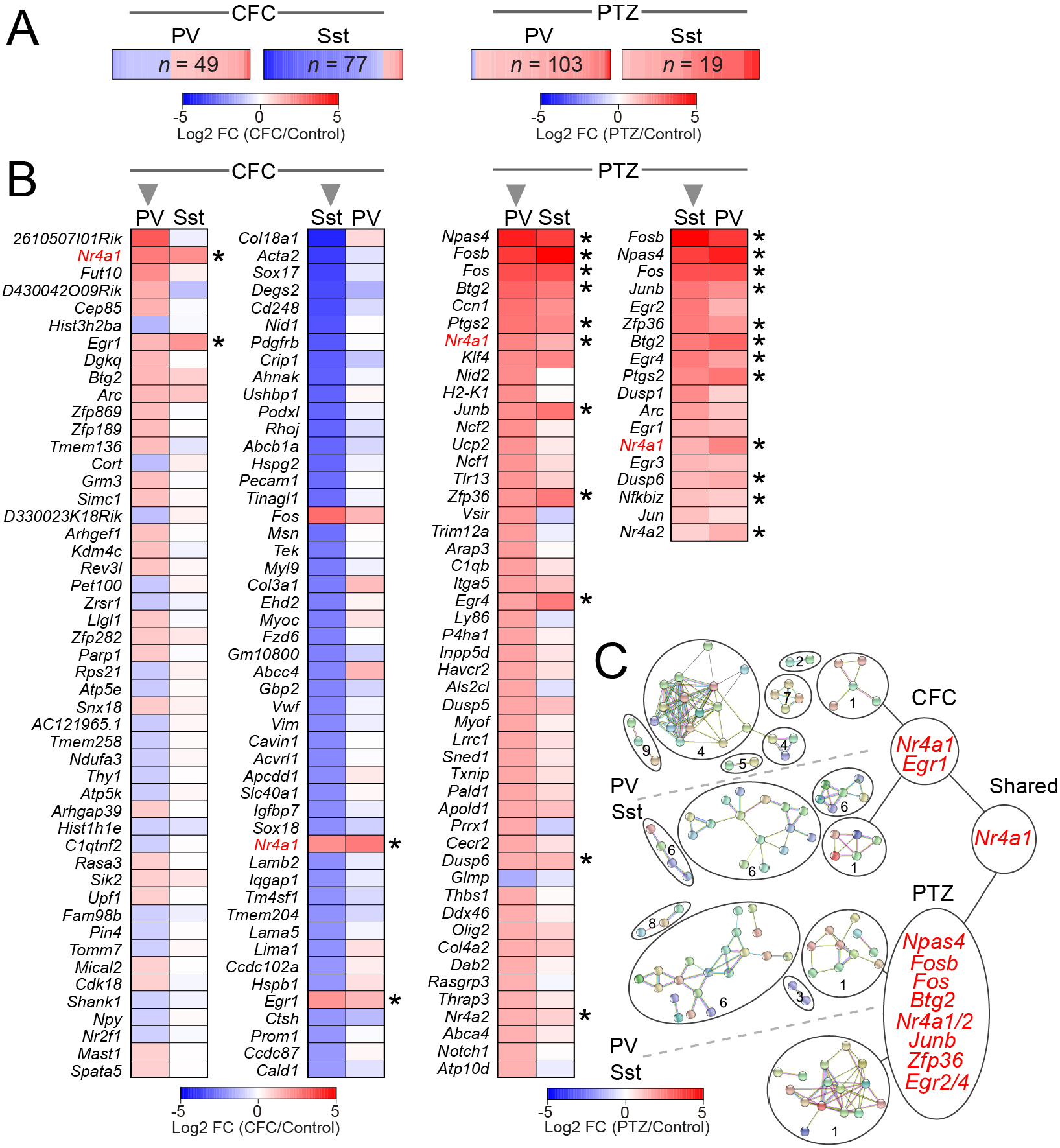
RNA-seq analysis of activity-dependent transcription in PV^+^ and Sst^+^ INs of p60 mice. **(A)** Heatmaps of mRNA levels of all differentially expressed genes in cortical/hippocampal libraries from *Pvalb*^*Cre*^/*RiboTag* and *Sst*^*Cre*^/*RiboTag* mice subjected to contextual fear conditioning (30 minutes post-CFC) or treated with PTZ (i.p., 50 µg/gm, 30 minutes post-injection). *n* = 3 to 4 mice/group. **(B)** Pairwise comparison of top hits. Heatmaps for significantly up- and down-regulated genes are ranked by fold change of mRNA levels in each IN population (arrows). Asterisks mark genes whose expression was significantly altered in both populations. Two middle columns show partial datasets. **(C)** STRING pathway analysis of gene functions (disconnected nodes not displayed). Circles mark the following proteins and/or pathways: 1 = Transcription factors (shared sets of induced TFs are shown in red); 2 = Ribosomal; 3 = mRNA splicing; 4 = Mitochondrial; 5 = Ubiquitination; 6 = Cytoskeleton and extracellular matrix; 7 = Neuropeptide signaling; 8 = Notch signaling; 9 = DNA repair.

**Fig. S3.**
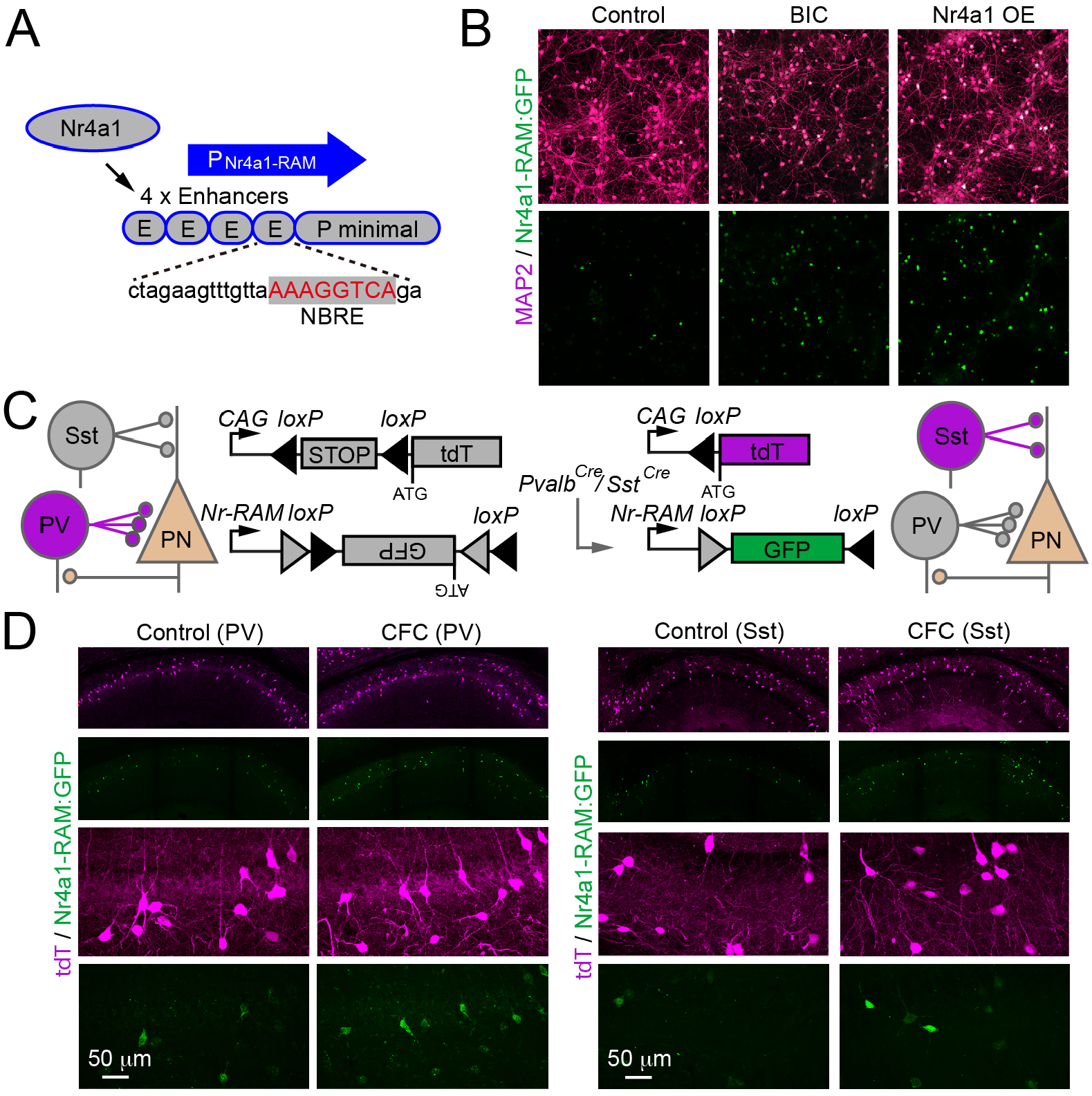
Analysis of transcriptional activity of Nr4a1. **(A)** Schematics of Nr4a1-RAM reporter. **(B)** Induction of Nr4a1-RAM:GFP in DIV14 primary cortical cultures under the normal conditions, 24 hours after application of the GABA receptor blocker, Bicuculine (BIC, 50 mm), or in the presence of exogenous Nr4a1 (OE). Nr4a1-RAM:GFP and full-length wildtype Nr4a1 were introduced with AAVs. **(C)** Schematics of Cre-dependent expression of Nr4a1-RAM:DIO-GFP and tdTomato (tdT, encoded by the *Ai9* allele) in PV^+^ and Sst^+^ INs. **(D)** Nr4a1-RAM:DIO-GFP was introduced into the CA1 of *Pvalb*^*Cre*^/*Ai9* and *Sst*^*Cre*^/*Ai9* mice with an AAV and monitored in p60 home-caged animals (Control) or 24 hours after CFC. Typical confocal images of coronal sections are shown.

**Fig. S4.**
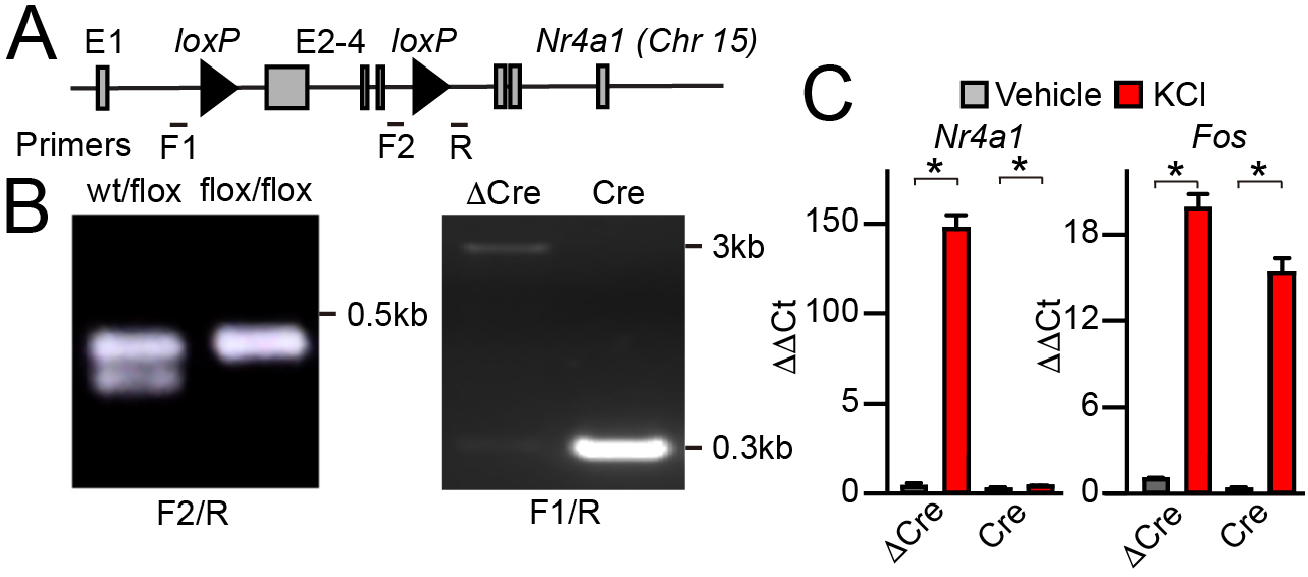
Validation of *Nr4a1*^*loxP/loxP*^ allele. **(A)** Schematics of *Nr4a1* ^*loxP/loxP*^ allele. LoxP sites flanking exons 2-4 and positions of genotyping primers are shown. **(B)** Left: genotyping of tail DNA samples from *Nr4a1*^*wt/loxP*^ and *Nr4a1*^*loxP/loxP*^ mice. Right: genotyping of mixed cortical cultures from *Nr4a1*^*loxP/loxP*^ mice. Cultures were infected with LVs expressing inactive (Δ) or active Cre recombinase under the control of the Ubiquitin promoter. **(C)** qPCR analysis of *Nr4a1* (left) and *Fos* (right) mRNA levels in cultures from *Nr4a1*^*loxP/loxP*^ mice carrying ΔCre or Cre, as expressed from LVs. Samples were collected under the normal conditions or after 30 minute depolarization with KCl (50 mM). Note lack of induction of Nr4a1 in the presence of Cre. Data from 3 independent experiments are plotted as Mean±S.E.M.

**Fig. S5.**
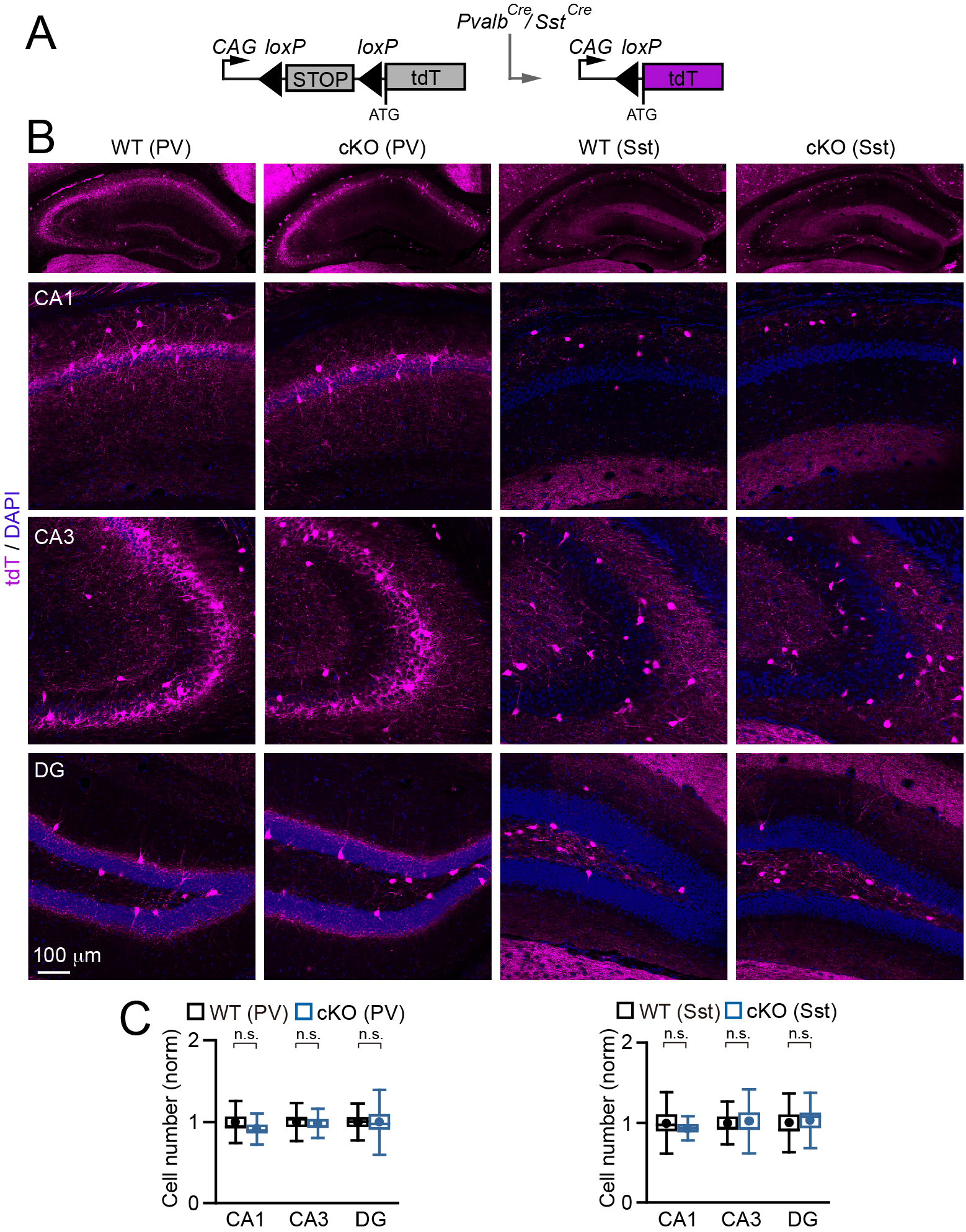
Preserved distribution and survival of INs in *Pvalb*^*Cre*^*/Nr4a1*^*loxP/loxP*^*/Ai9* and *Sst*^*Cre*^*/Nr4a1*^*loxP/loxP*^*/Ai9* cKO mice. **(A)** Schematics of Cre-dependent expression of tdTomato (tdT) from the *Ai9* reporter allele. **(B)** Representative confocal images of labeled PV^+^ and Sst^+^ cells in the hippocampi of p60 mice. **(C)** Quantifications of IN numbers in the CA1, CA3 and dentate gyrus (DG). Graphs represent mean values (circles), standard errors (boxes), standard deviations (whiskers), and medians (horizontal lines). cKO values were normalized to WT littermates in each pair of mice. PV^+^ (WT), *n* = 4 mice/24 sections; PV^+^ (cKO), *n* = 4/24; Sst^+^ (WT), *n* = 3/18; Sst^+^ (cKO), *n* = 3/18. *p* values for WT vs cKO were defined by t-test: PV^+^ (CA1), *p* = 0.2; PV^+^ (CA3), *p* = 0.8; PV^+^ (DG), *p* = 0.99; Sst^+^ (CA1), *p* = 0.48; Sst ^+^ (CA3), *p* = 0.87; Sst ^+^ (DG), *p* = 0.8.

**Fig. S6.**
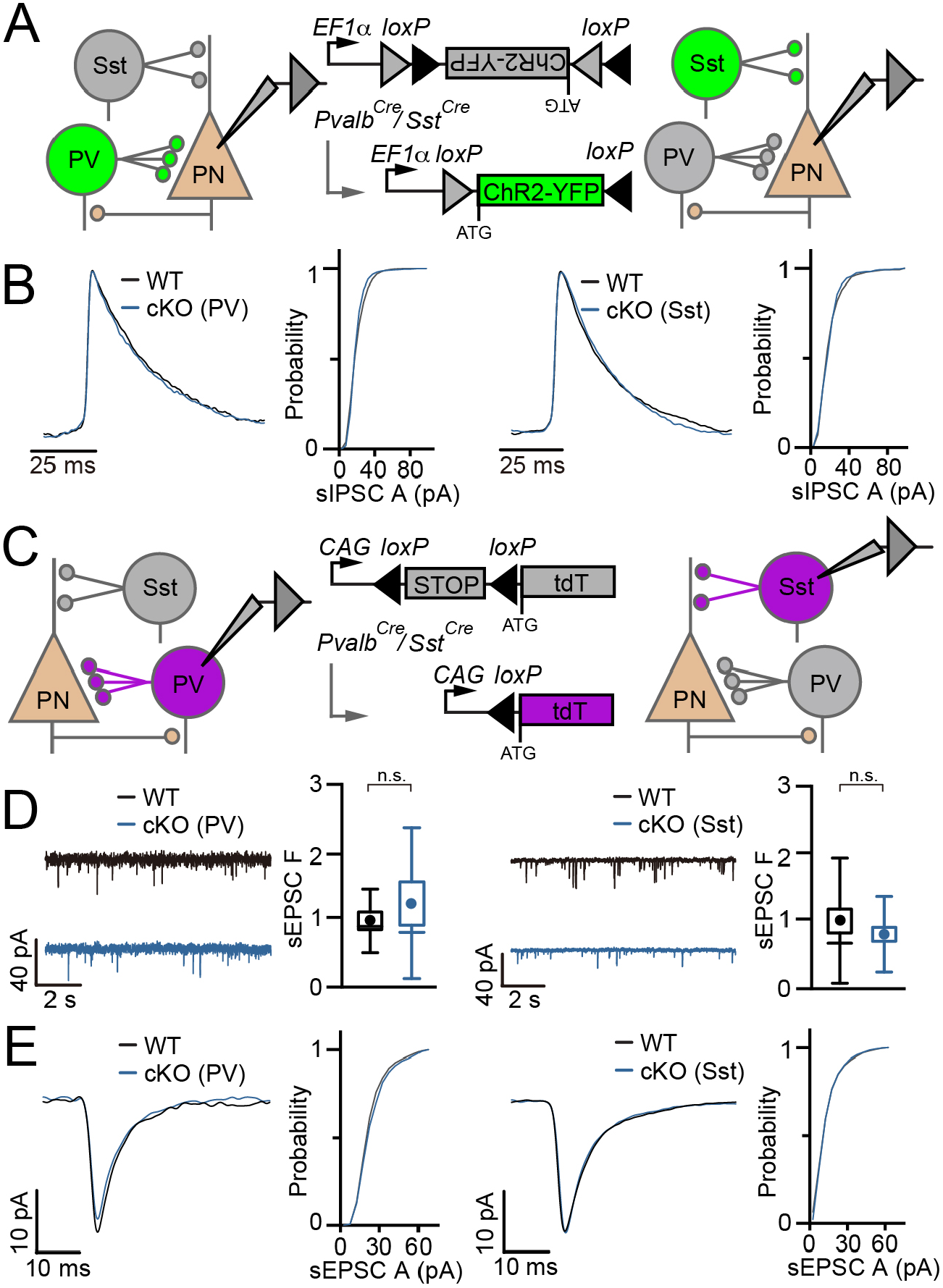
Extended electrophysiological analysis of synaptic transmission. **(A** and **B)** Spontaneous inhibitory postsynaptic currents (IPSCs) were recorded from CA1 PNs in acute brain slices from Nr4a1 WT (*Nr4a1*^*wt/wt*^) and IN-specific cKO mice (*Nr4a1*^*loxP/loxP*^) carrying corresponding Cre drivers and Cre-dependent ChR2-YFP, as expressed from an AAV. Panels show recording configurations (A), sample traces of quantal sIPSCs, and cumulative probability histograms of sIPSC amplitudes (B). PV^+^ (WT), *n* = 6 mice/25 neurons; PV^+^ (cKO), *n* = 6/21; Sst^+^ (WT), *n* = 3/22; Sst^+^ (cKO), *n* = 3/25. **(C** to **E)** Whole-cell recordings from INs of Nr4a1 WT and cKO mice carrying the *Ai9* tdTomato reporter. **(C)** Schematics of IN labeling and recording configurations. **(D)** Sample traces of spontaneous excitatory postsynaptic currents (sEPSCs) and quantifications of sEPSC frequencies. Graphs represent mean values (circles), standard errors (boxes), standard deviations (whiskers), and medians (horizontal lines). cKO values were normalized to WT littermates in each pair of mice. *p* values for WT vs cKO were defined by t-test: PV^+^, *p* = 0.47; Sst^+^, *p* = 0.35. **(E)** Sample traces of quantal sEPSCs and cumulative probability histograms of sEPSC amplitudes. PV^+^ (WT), *n* = 4 mice/13 neurons; PV^+^ (cKO), *n* = 4/12; Sst^+^ (WT), *n* = 3/27; Sst^+^ (*Nr4a1* cKO), *n* = 3/30.

**Fig. S7.**
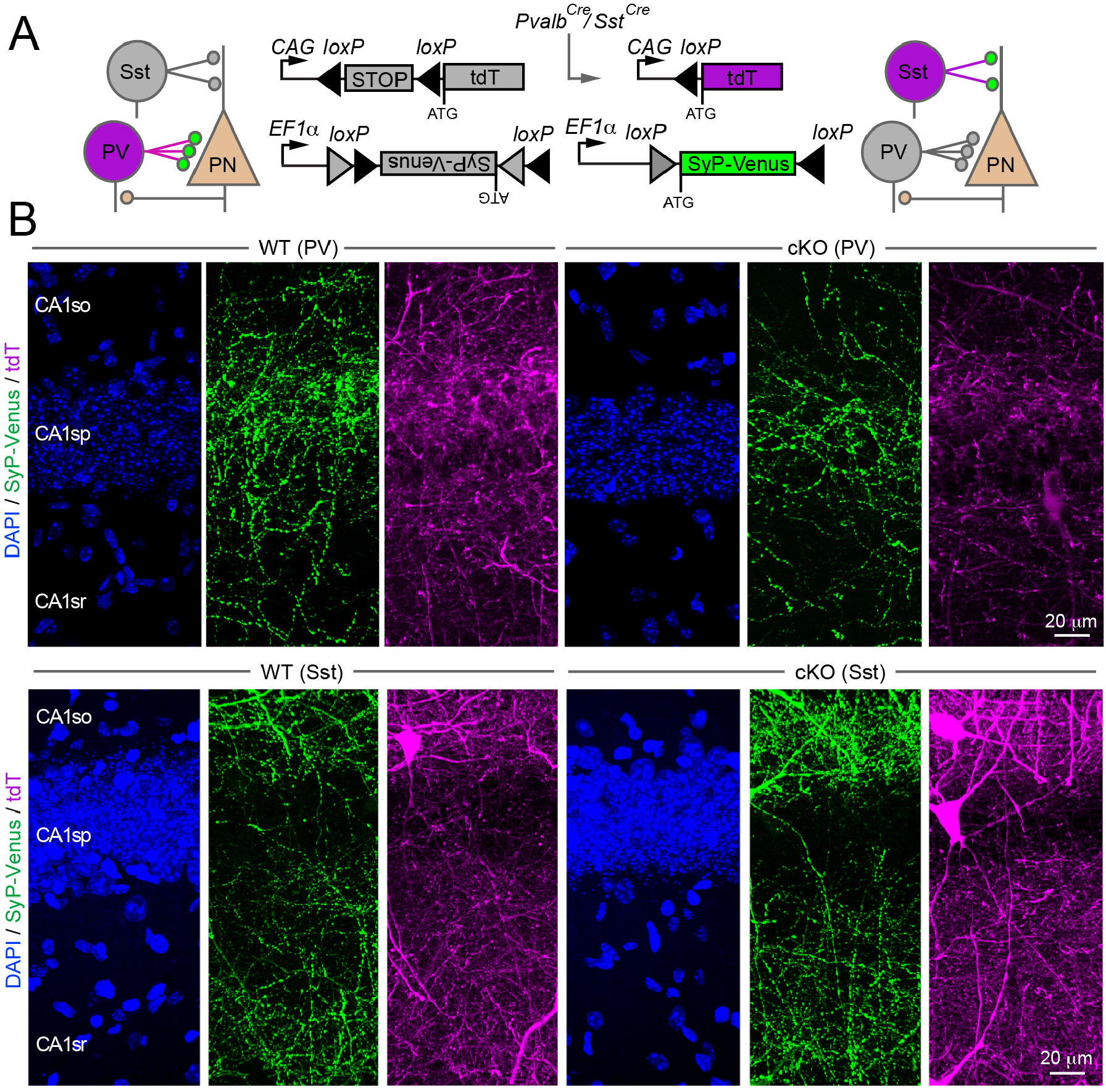
Preserved axonal targeting of Nr4a1-deficient INs. **(A)** Schematics of dual Cre-dependent labeling of PV^+^ and Sst^+^ cells in Nr4a1 WT and cKO mice with the *Ai9* tdTomato (tdT) reporter and virally expressed presynaptic marker, Synaptophysin (SyP)-Venus. **(B)** Representative confocal images of labeled projections and SyP-Venus-positive nerve terminals in the CA1 at p60. CA1so = stratum oriens; CA1sp = pyramidal cell layer; CA1sr = stratum radiatum. Note preserved segregation of terminals along the CA1so-CA1sp-CA1sr axes in both cKO lines.

**Fig. S8.**
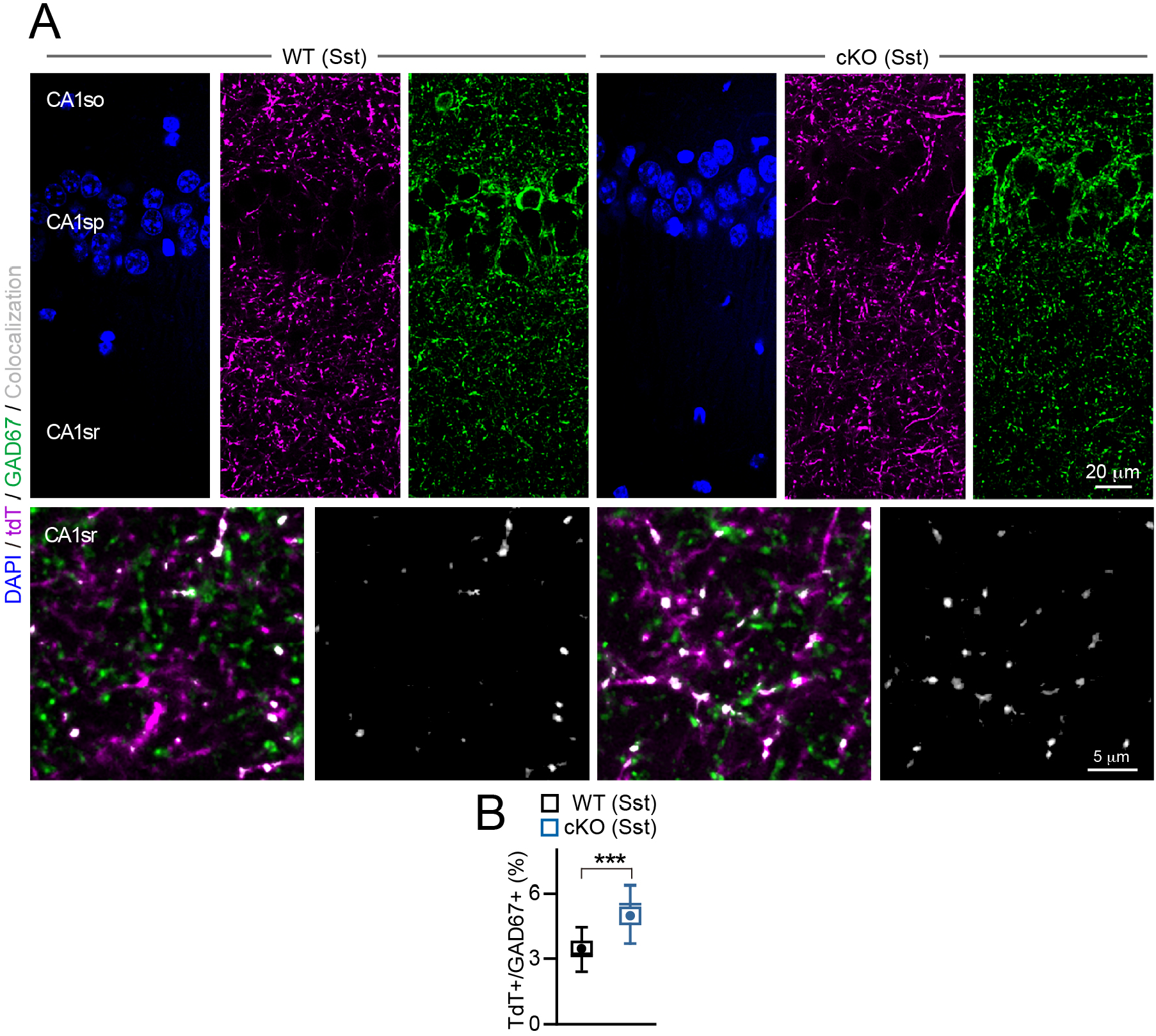
Distribution of GABAergic terminals in Nr4a1 WT and Sst^+^ IN-specific cKO mice. **(A)** Confocal images of the CA1 in animals carrying the *Ai9* tdTomato (tdT) reporter. Brain sections from p60 animals were labeled with the antibody against pan-inhibitory presynaptic marker, GAD67. **(B)** Co-localization of GAD67-positive boutons with tdTomato. Graph represents mean values (circles), standard errors (boxes), standard deviations (whiskers), and medians (horizontal lines). *n* = 3 mice/genotype, *p* = 0.004 (defined by t-test).

**Fig. S9.**
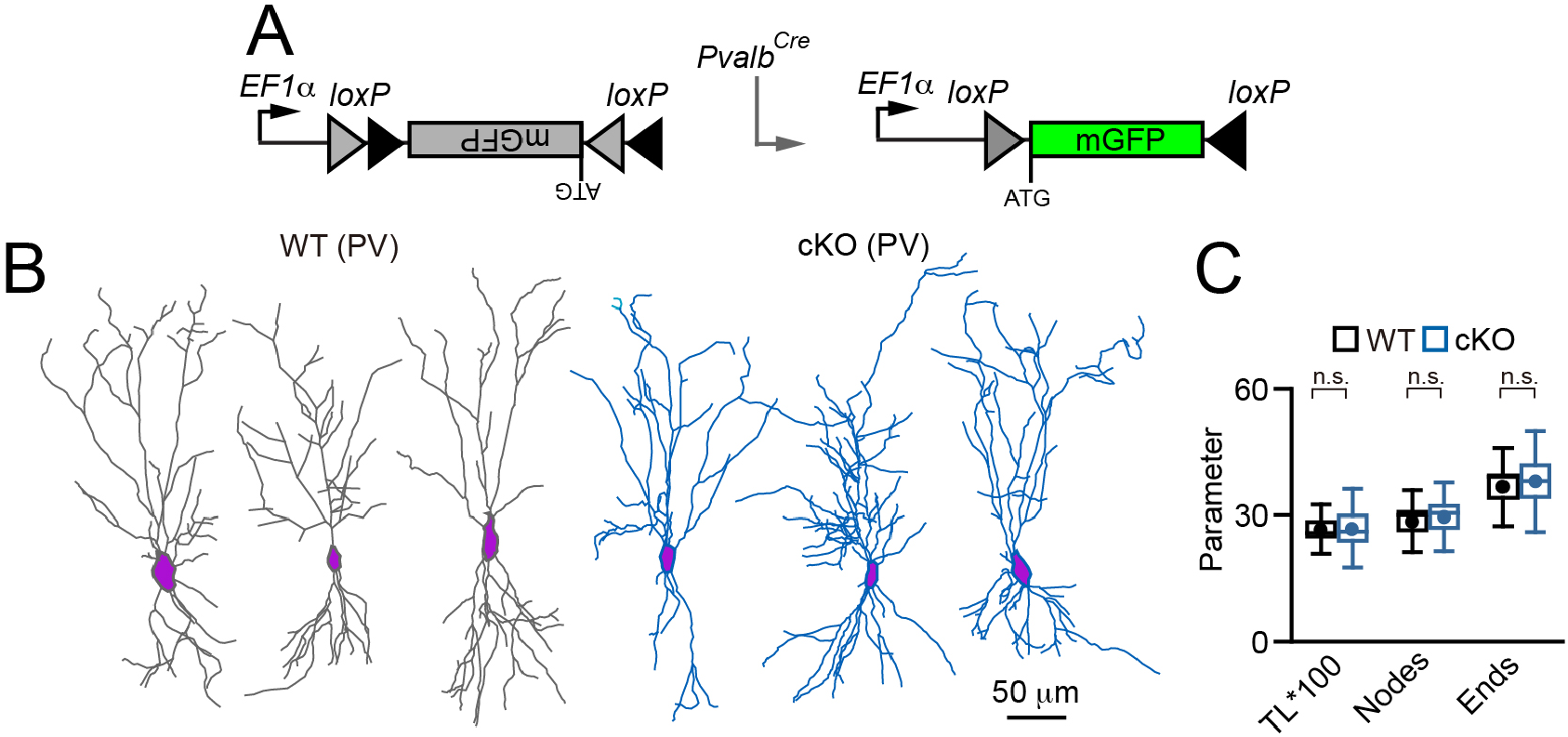
Dendrite morphologies of normal and Nr4a1-deficient PV^+^ cells. **(A)** Schematics of Cre-dependent expression of mGFP reporter from an AAV. **(B)** Reconstructions of dendritic trees of individual INs labeled with mGFP in p60 Nr4a1 WT and cKO mice **(C)** Quantifications of total dendrite lengths (TL) and numbers of nodes and ends. Graph represents mean values (circles), standard errors (boxes), standard deviations (whiskers), and medians (horizontal lines). *n* = 3 mice/10 to 12 neurons per genotype). TL, *p* = 0.94; Nodes, *p* = 0.76; Ends, *p* = 0.78 (defined by t-test).

**Fig. S10.**
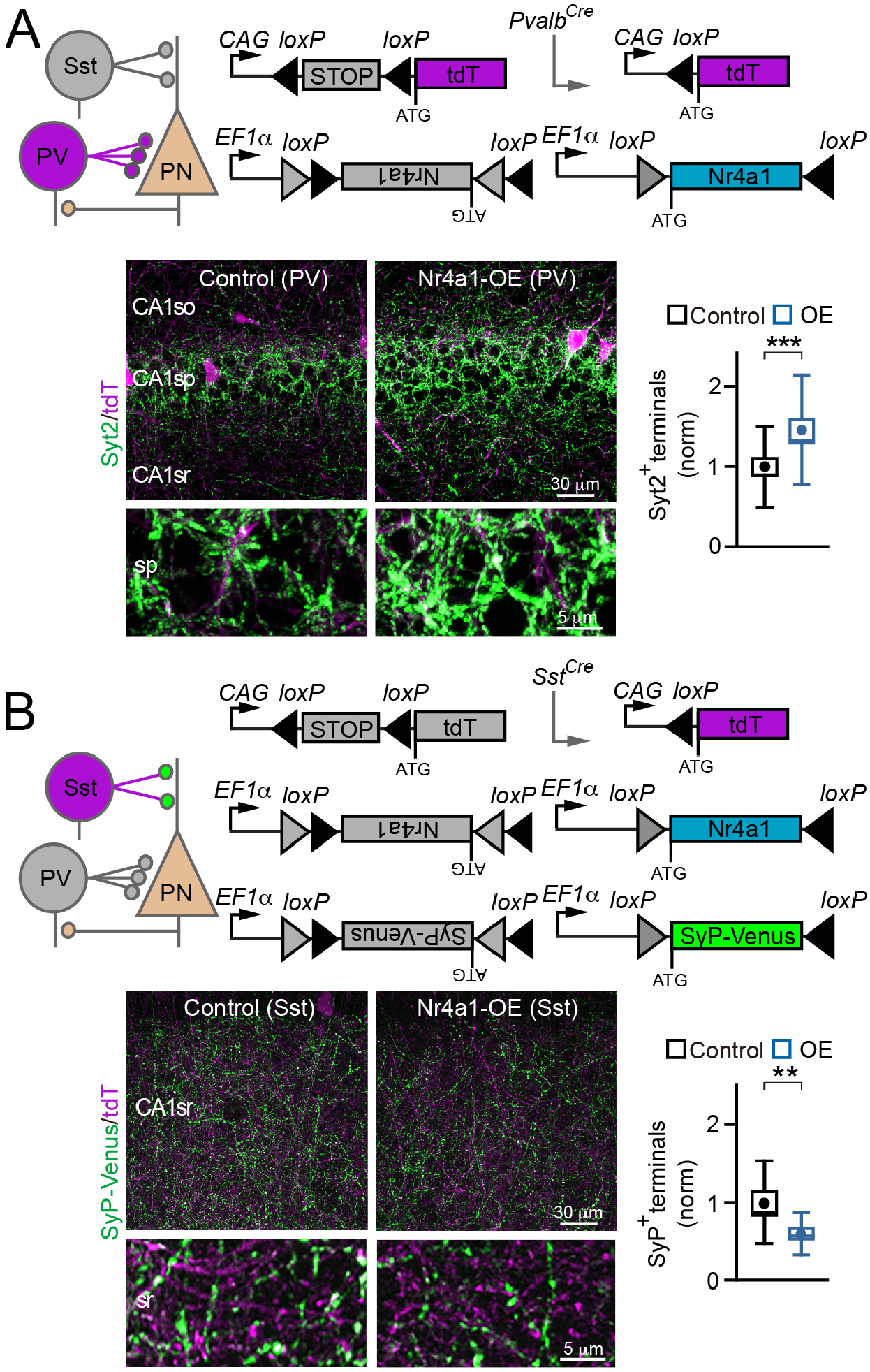
Effects of Nr4a1 gain-of-function on presynaptic networks of PV^+^ and Sst^+^ INs. **(A)**The full-length Nr4a1 cDNA was overexpressed (OE) from a Cre-dependent AAV in PV^+^ cells of neonatal *Pvalb*^*Cre*^*/Ai9* mice. Panels show schematics of Nr4a1 and reporter induction, typical confocal images of brain sections stained with the antibody against the maker of presynaptic terminals of PV^+^ cells, Syt2, and quantifications of somatic Syt2-positive puncta in CA1sp. Control, *n* = 4 mice/24 sections; OE, *n* = 4/27. *p* = 0.008 (defined by t-test). **(B)**The full-length wildtype Nr4a1 cDNA and the presynaptic marker, SyP-Venus, were introduced with Cre-dependent AAVs into Sst^+^ cells of neonatal *Sst*^*Cre*^*/Ai9* mice. Panels show schematics of Nr4a1 and reporter induction, typical confocal images of brain sections, and quantifications of dendritic SyP-Venus-positive puncta in CA1sr. Control, *n* = 3/16; OE, *n* = 3/16. *p* = 0.011 (t-test). In both panels, graph represents mean values (circles), standard errors (boxes), standard deviations (whiskers), and medians (horizontal lines). All data were collected at p60. OE values were normalized to control littermates in each pair of mice.

**Fig. S11.**
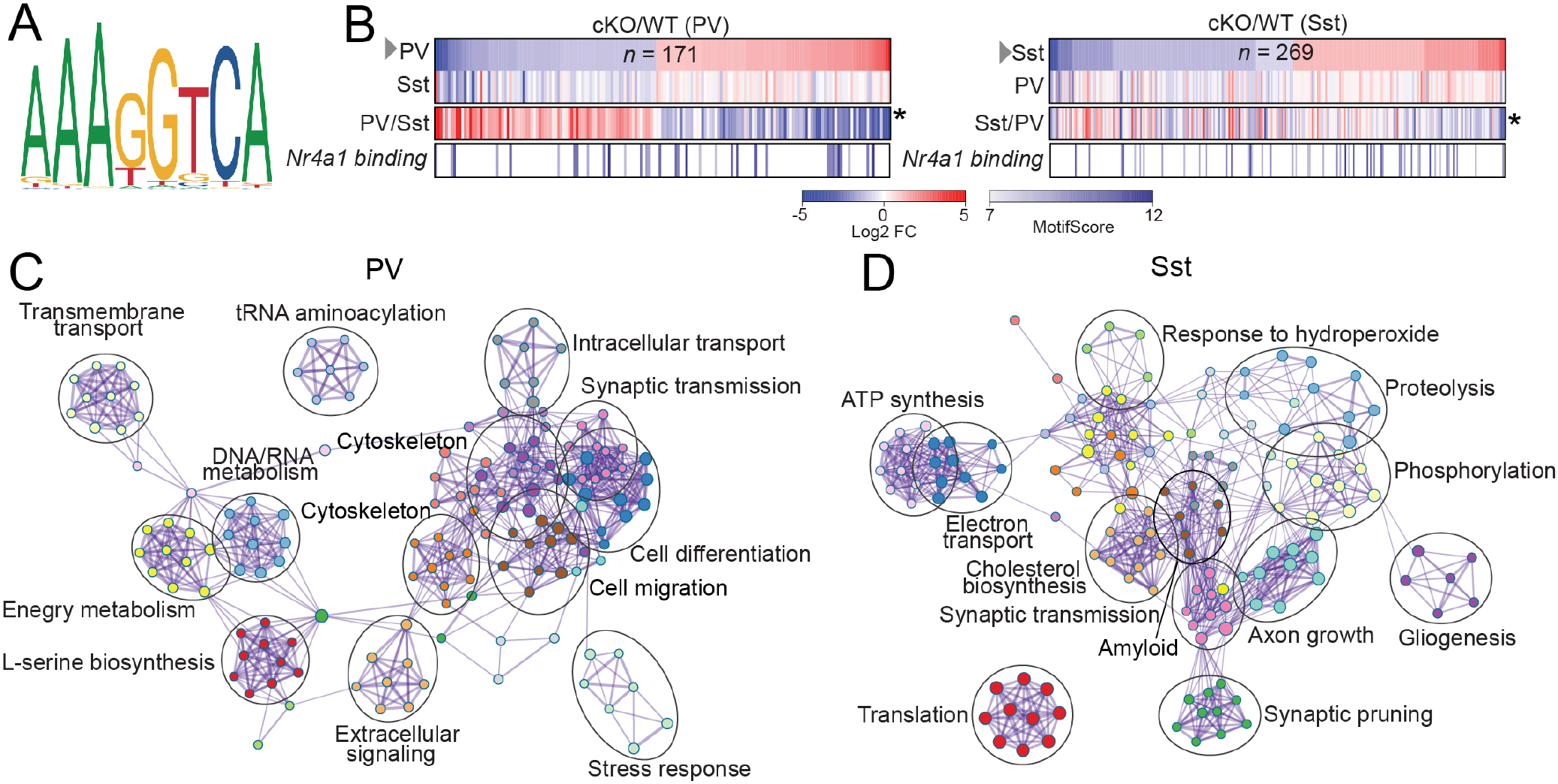
RNA-seq analysis of gene expression in Nr4a1-deficient PV^+^ and Sst^+^ INs. **(A)** Nr4a1 binding motif. **(B)** Heat maps show comparisons of mRNA levels of genes that were significantly up- or down-regulated in each population of Nr4a1-deficient INs (top rows, -1<log2 FC>1), with their expression in other Nr4a1-deficient population (second rows) and relative expression in Nr4a1 WT mice (third rows). Bottom rows demonstrate the scores for Nr4a1 binding motif, as defined by Homer v4.11. **(C** and **D)** Pathway analyses for genes that are mis-regulated in Nr4a1-deficient PV^+^ and Sst^+^ cells.

**Fig. S12.**
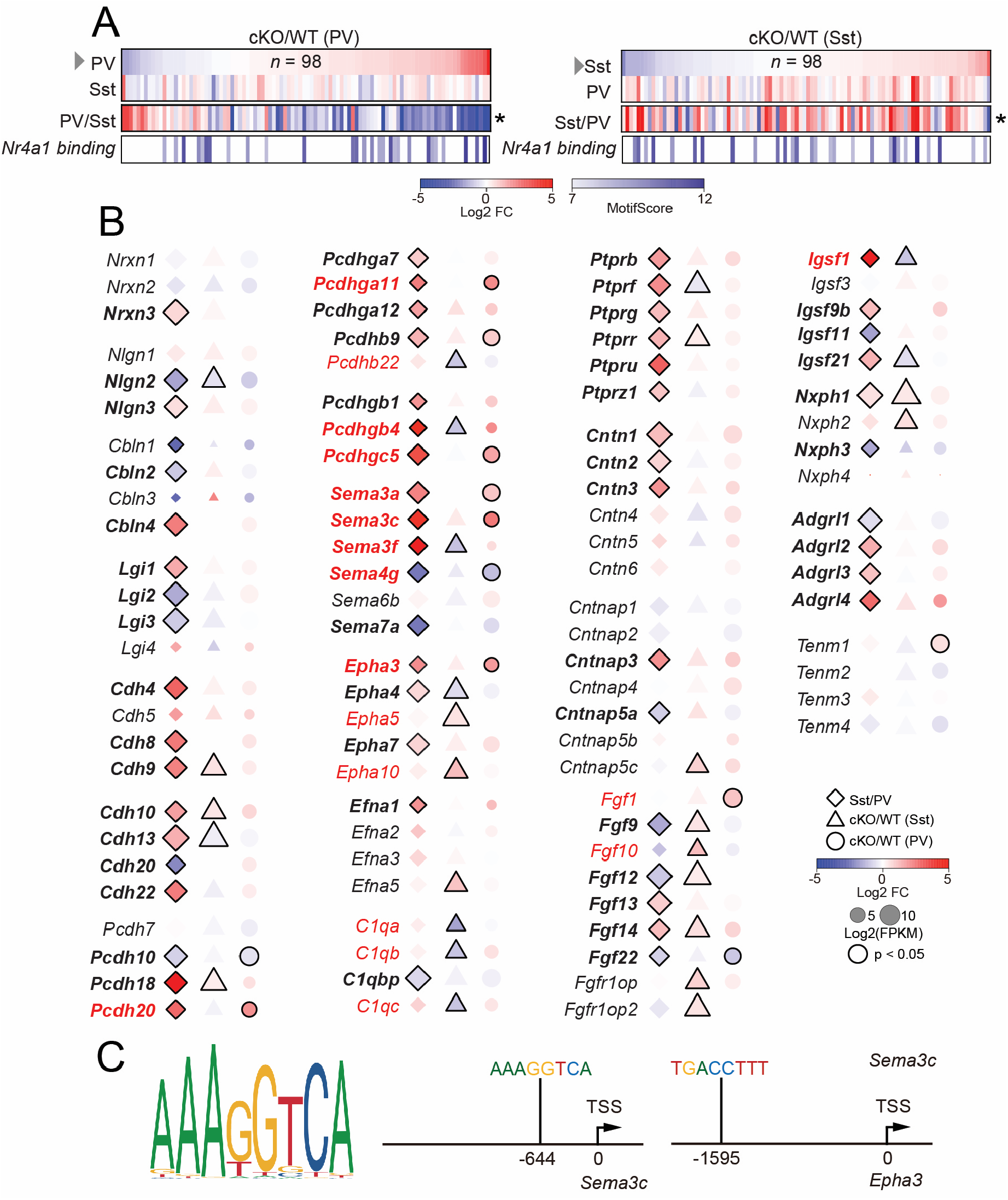
Expression of gene families essential for synaptic differentiation in normal and Nr4a1-deficient INs. **(A)** Heatmaps of mRNA levels of 98 genes involved in synaptic differentiation are ranked by fold-change in each population of Nr4a1-deficient INs. Panels also show the expression of same genes in other Nr4a1-deficient population (second rows), their relative expression in Nr4a1 WT mice (third rows) and the scores for Nr4a1 binding motif, as defined by Homer v4.11 (bottom rows). **(B)** Annotated heatmaps of mRNA levels of individual genes demonstrate their relative expression levels in normal INs (first columns, plotted as Sst/PV ratios) and expression in each population of Nr4a1-deficient INs (second and third columns). **(C)** Positions of Nr4a1 binding motifs upstream transcription start sites (TSS) in *Sema3c* and *Epha3* genes.

